# Mycorrhizal symbiont provides growth benefits in host plants via phosphate and phenylpropanoid metabolism

**DOI:** 10.1101/2023.07.06.547905

**Authors:** Cheng-Yen Chen, Naweed I. Naqvi

## Abstract

Using functional interaction assays, *Tinctoporellus species* isolate AR8 was identified as a plant growth-promoting fungus from Arabidopsis roots.

Confocal microscopy revealed interstitial growth and intracellular endophytic colonization within root cortex by AR8 hyphae prior to induction of beneficial effects.

AR8 improved plant growth and fitness across a broad range of monocot and dicot host species. AR8 solubilized inorganic phosphate and enabled macronutrient phosphorus assimilation into the host plants, and the resultant growth promotion required an intact phosphate starvation response therein.

Metabolomics analysis identified a highly specific subset of primary and secondary metabolites such as sugars, organic acids, sugar alcohols, amino acids, and phenylpropanoids, which were found to be essential for the plant growth-promoting activities of AR8.

*trans*-Cinnamic acid was identified as a novel AR8-induced plant growth promoting metabolite.

## Introduction

Plants provide ecological niches for a plethora of microorganisms, which build highly complex communities therein, and profoundly influence the overall functioning and stability of such tripartite plant-microbe-environment ecosystems. Numerous studies have revealed that such microbial associations affect seed germination, growth and development, plant nutrition, resilience, and reproduction in the plant hosts (Berendsen et al., 2012; Vandenkoornhuyse et al., 2015; Leach et al., 2017). Given these paramount functions, such microbial assemblies are seen as an extended or second plant genome, and are also referred to as microbiota (comprising all microorganisms) or microbiome (comprising all microbial genomes). The rhizosphere is a ubiquitous region for a set of microbial populations that support plant fitness too (Lundberg et al., 2012; Ofek-Lalzar et al., 2014). The below-ground interactions between roots and microbiota are critical for the plant adaptation capacity, which can determine plant growth and health through various direct or indirect mechanisms (Bever et al., 2012; Herrera Paredes and Lebeis, 2016). Therefore, it is of major interest to study plant microbiota that enhance growth, biomass and/or provide resistance against pathogen infection in the host plants.

The mycobiome, or mycobiota, is the fungal community associated with an organism or a habitat. Dependent on the multiple pivotal ecological functions driven by soil fungi, plant performance can be governed and modified by changing the rhizosphere mycobiota. Arbuscular mycorrhizal fungi (AMF) are a characteristic example of plant growth-promoting fungi (PGPF) in the symbiotic system. It is now widely accepted that AMF symbiosis contributes to plant nutrition (mainly phosphate; Pi), including solubilization, mineralization, and/or transfer of nutrients in the bioavailable forms, which allows plants to thrive in harsh environments with poor nutrient availability (Hijikata et al., 2010; Balliu et al., 2015; Etesami et al., 2021). The role of root fungal endophytes in beneficial interactions with plants has been underestimated but has recently attracted a great deal of attention due to their ability to provide mycorrhizal-like functions to non-mycorrhizal plants. For example, *Colletotrichum tofieldiae* solubilizes plant-inaccessible hydroxyapatite and transfers Pi to *Arabidopsis thaliana* under low-Pi conditions (Hiruma et al., 2016). Similarly, the Helotiales fungus F229 transfers Pi to *Arabis alpina* under both low-and high-Pi conditions (Almario et al., 2017). These studies endorse that fungus-to-plant Pi transfer, commonly assigned as a hallmark of mycorrhizal symbiosis, also occurs in root fungal endophytes and non-mycorrhizal plant interactions. Given this scenario, in-depth studies of symbiotic mycobiota members and their function(s) on non-mycorrhizal plants are critical to expand our knowledge of the ecological relevance of these associations for plant growth and development.

The advantage of root fungal endophytes for plant growth promotion through different beneficial pathways shows great potential in developing new strategies for sustainable agriculture (Calvo Velez et al., 2014; Olanrewaju et al., 2017; Sood et al., 2020). Despite their benefits, the use of root fungal endophytes in agriculture remains far below the number of fungi described with plant growth-promoting activities. Lack of knowledge about molecular features of plant-fungus symbiosis is the main issue limiting the use of these PGPF as sustainable alternatives. Metabolomics analysis focusing on plant-fungal interaction has shown that the specialized metabolic phenotypes of host plants are essential for shaping the morpho-physiological traits of functional symbiosis with AMF. For example, the carbon demands of AMF affect the photosynthetic capacity of plants, thereby enhancing sugar accumulation and increasing plant biomass (Kaur and Suseela, 2020; Kaur et al., 2022). Furthermore, the reprogramming of secondary metabolism in carotenoid, flavonoid, and phenylpropanoid biosynthesis due to AMF symbiosis enables the plants to better stress tolerance (Fester et al., 2005; Scervino et al., 2005; Schweiger and Müller, 2015; da Silva and Maia, 2018). Thus, the specificity of plant metabolome not only illustrates molecular signatures but also reflects symbiotic outcomes in the plant-fungal interactions. More importantly, understanding the dynamics of plant metabolites in symbiotic associations would help decipher the plant growth-promoting mechanisms, especially since the functional symbiosis elicited by root fungal endophytes is highly host-specific.

Plants of the Brassicaceae family are found within the group of non-mycorrhizal symbiosis, which comprises several model species of great scientific interest (i.e., *A. thaliana*) and/or agronomic importance (i.e., food crops within the genus *Brassica*) (Cosme et al., 2018). On the basis of accumulating evidence, the interactions between root fungal endophytes with plant hosts resemble mycorrhizal symbiosis. However, most studies were conducted on the model plants in controlled and optimized growth environments while only a few were performed under less favorable field conditions. In this study, the primary objectives were to harness mycobiome-based functions to improve plant growth by: (1) the identification of root fungal endophyte(s) that can benefit green leafy vegetable Choy Sum (*Brassica rapa var. parachinensis*) (2) the analysis of plant metabolome underlying the beneficial effects imparted by the root fungal endophyte to the host plant; (3) the field trials to clarify plant growth-promoting activity of root fungal endophyte and its ecological relevance as a potential biofertilizer for agriculture.

Here, we identify a novel mycorrhizal fungus, the *Tinctoporellus* species isolate AR8 (hereafter AR8), which demonstrated robust plant growth-promoting activity in green leafy vegetables in indoor and field conditions. Characterization of AR8-inoculated Choy Sum revealed that AR8 hyphae systemically colonized the inter-and intracellular spaces within the host root cortex and thus underscored the capability of fungal hyphae to transfer soil nutrients to host plants. Indeed, *in vitro* studies confirmed that the contribution of AR8 to plant growth promotion involves the solubilization of inorganic Pi and the transfer of bioavailable phosphorus by root-associated hyphae. Detailed metabolite profiling extended our findings by providing mechanistic and functional insight into the repertoire of primary and secondary metabolites that significantly changed in Choy Sum upon AR8 inoculation. We also identified and characterized *trans*-cinnamic acid (*t*-CA) as the metabolite with plant growth-promoting activity. Increased shoot biomass by *t*-CA in exogenous complementation assays provided further evidence for its bioactive functions, likely associated with enhanced plant growth in the AR8 symbiosis model. Collectively, our results support a mycorrhizal model system that imparts beneficial functions in plant growth via a new member of rhizosphere mycobiota with potential applications in improving productivity in traditional and modern urban farm crops.

## Materials and Methods

### Plant materials and growth conditions

*Arabidopsis thaliana* ecotype Columbia (Col-0) wild type, the mutant line *phr1*, Choy Sum (*Brassica rapa var. parachinensis*), Kailan (*Brassica oleracea var. alboglabra*), rice cultivar CO39, and barley cultivar Express were used in this study. Seeds were stored at 4°C in the dark to allow stratification. Seeds were surface-sterilized with 70% ethanol for 5 min, 10% commercial bleach for 2 min, and followed by five times rinsing in sterile distilled water. Sterilized seeds were placed on full Murashige and Skoog basal salts medium (Sigma-Aldrich; no. M5524) with 1% sucrose and 1% agar (MS-agar) for germination and incubated vertically in a plant growth chamber (AR95L, Percival Scientific) under long day conditions (16 h /8 h light/dark photoperiod) at 22°C with 60% relative humidity and 120 μmol m^-2^ s^-1^ light intensity. For plant growth promotion assays, seedlings were transferred to trays containing autoclaved (121°C, 1 hour) peat/perlite mix (BVB Substrates, Netherlands) prior to fungal inoculation. Plants in the tray were watered (1000 ml/day) according to the method described previously (Tan et al., 2020), and no additional fertilizers were added during the experimental period.

### Plant growth promotion assays

For qualitative and quantitative analysis of the growth-promoting effects of fungal isolates extracted from Arabidopsis rhizosphere, conidial suspension (10^6^ spores) were inoculated into the rhizosphere region of Choy Sum plants. Four day-old Choy Sum seedlings were transferred to the sterilized soil and conidia were directly inoculated to the roots and surrounding soil. Distilled water was used as a mock control with the same inoculation method. Shoot fresh weight was measured at 7, 14, and 21 days post-inoculation (dpi). To further verify the effect of the fungal isolate AR8 on reproductive growth, the number of siliques was counted at 49 dpi. *A. thaliana* Col-0, Kailan, rice, and barley were used as models for monocot and dicot species to test AR8 growth-promoting effect using the same settings and methodology.

### Confocal microscopy and imaging

For microscopic analysis of fungus-root interactions, four day-old Choy Sum seedlings were inoculated with AR8 conidia (10^6^ spores) for 16 and 24 hours post-inoculation (hpi) (conidia morphology and germination), and 4, 7, 14 dpi (root colonization). Distilled water was used as a mock control with the same inoculation method. Conidia and roots were stained with 10 μg/ml Wheat Germ Agglutinin, Alexa Fluor™ 488 Conjugate for 10 minutes and 10 μg/ml Propidium iodide for 10 minutes to visualize fungal structure and root cell walls, respectively. Imaging was done on a Leica TCS SP8 X inverted confocal system equipped with an HC Plan Apochromat 20×/0.75 CS2 Dry objective or a 63×/1.40 CS2 Oil objective. Green fluorescence (Wheat Germ Agglutinin Alexa Fluor™ 488 Conjugate) was excited at 488 nm and detected at 500-530 nm and Red fluorescence (Propidium iodide) excited at 561 nm and detected at 600- 700 nm. All parts of the system were under the control of Leica Application Suite X software package (release version 3.5.5.19976).

### Elemental analysis by Inductively-Coupled Plasma mass spectrometry (ICP-MS)

Samples of 21 dpi Choy Sum shoot (with or without AR8 inoculation) were dried for 5 days at 65 °C. For pre-digestion, Homogenized plant powder (100 mg) was performed by 2.5 ml of HNO_3_ (66% v/v) and 0.5 ml of H_2_O_2_ (30% v/v) overnight. High Performance Microwave Digestion System was followed to digest samples for 3 hours. Final solution was diluted by 1:10 dilution with deionized water and stored at 4 °C before analysis. The determination of nitrogen, phosphorus, and potassium levels was performed with an Agilent 7700 ICP-MS (Agilent) following the manufacturer’s instructions.

### Pi translocation assay

Square petri dishes (root/hyphal compartment, RHC; 12.5 x 12.5 cm) were prepared with either Pi-limiting (100 μM KH_2_PO_4_) or Pi-rich (1250 μM KH_2_PO_4_) sucrose-free MS-agar medium. Inside each square petri plate, a circular petri dish (hyphal compartment, HC; 3.8 cm in diameter) was prepared with Pi-rich sucrose-free MS-agar medium and placed at the bottom. Notably, the two compartments were separated by the plastic wall of the circular petri dish, which impeded the growth of plant roots towards the content of the smaller plate. Two PA agar plugs (5 mm) with or without AR8 mycelia were inoculated to HC for pre-inoculation 7 days. During the pre-inoculation, AR8 hyphae spread from HC to the RHC, bridging the two compartments. Seven day-old *A. thaliana* Col-0 or *phr1* mutant seedlings were transferred to the RHC and cultivated vertically for 7 days. Shoot fresh weight was measured to determine the effect of fungal Pi transport activity.

### Metabolomics profiling

Metabolite profiling was performed by gas chromatography coupled to electron impact ionization/time-of-flight mass spectrometry (GC-EI/TOF-MS) using Agilent 7890A gas chromatogram with split and split-less injection onto the Agilent J&W GC column DB-5MS (30 m length, 0.25 mm inner diameter, 0.25 μm film thickness, Agilent), which was connected to a 7200 quadrupole time-of-flight mass spectrometer. Helium was used as carrier gas and the flow rate was 1 ml/min. Injection volume was 1 μl in split and split-less mode. Injection and transfer line temperatures were 250°C and 280°C, respectively. The oven temperature was held at 70°C for 1 minute, and then increased to 250°C at 10°C/min and then it was increased to 300°C at 25°C/min and held for 6 minutes. The GC total run time was 27 minutes. The solvent cut time was 4 minute. The ion source was operated in electron ionization (EI) mode and its temperature was 230°C. The scan range for TOF was from m/z 50 to 800.

### Statistical Analysis

Statistical analysis was carried out using GraphPad prism software (San Diego, CA. and the values of the treatments represented as mean with standard error. The significance of differences between the treatments was statistically evaluated using Student’s ‘t’ test and significance was considered at a probability level of *P* < 0.05 (*), *P* < 0.01(**), and *P* < 0.001 (***).

## Results

### Characterization of the mycobiota from *Arabidopsis thaliana* rhizosphere

We used extracts from the surface-sterilized roots of *A. thaliana* Col-0 to obtain the mycobiomes or fungal microbiota via subculture on PA medium. The fungal isolates thus obtained were taxonomically identified by amplifying and sequencing the ITS region of fungal the ribosomal DNA from the respective fungal isolates. Barcoding and identification of fungal DNA on the NCBI BLAST database (https://blast.ncbi.nlm.nih.gov/Blast.cgi) was subsequently conducted to assign the taxonomic identity of the mycobiont based on the highest ID scores. As a result, a total of 21 individual fungal genera, including 18 isolates from Ascomycota and 3 isolates from Basidiomycota, were obtained from *A. thaliana* roots. To further verify the impact (if any) on plant growth, we inoculated Choy Sum seedlings with our fungal isolates individually and cultivated them under soil conditions. Shoot fresh weight or biomass of the mock control and the fungus-inoculated plants was measured at 21 days (Fig. S1).

Based on our analysis, Choy Sum showed an increase in shoot biomass when co-cultivated individually with 9 of the 21 fungal isolates, thus suggesting their plant growth-promoting effect on Choy Sum under soil conditions (Fig. S1b). More importantly, the fungal isolate AR8 exhibited a better and more consistent beneficial effect on growth of Choy Sum shoots compared to the other PGPF isolates. Thus, we selected AR8 to further investigate such beneficial effects on plant biomass (Figure 1a,b). The significantly improved Choy Sum growth provided by AR8 under soil conditions was evident at 14 dpi. AR8-inoculated plants exhibited a significantly higher shoot biomass increase by 22.01% at 14 dpi, and 37.07% at 21 dpi (Fig. 1c,d S2a). To address the long-term impact of AR8 interaction/colonization, the reproductive growth of Choy Sum was also studied as an index of fertility. We observed that Choy Sum inoculated with the beneficial mycobiont AR8 displayed an earlier transition to flowering and an overall increase in production of siliques (Fig. S2b), thus indicating that the beneficial fungus AR8 promotes the growth and development during both the vegetative and reproductive stages in the host plants. We also tested the impact of AR8 on the model Brassicaceae species *A. thaliana* Col-0 and Kailan, and in the cereal crops rice and barley. Remarkably, not only Arabidopsis and Kailan but also barley demonstrated a significant improvement with an average increase of 53.61%, 22.85%, and 21.91%, respectively, upon AR8 inoculation under soil conditions (Fig. S3,S4c,d). However, AR8 showed a growth inhibitory effect on rice, and the shoot fresh weight of rice was significantly decreased by 18.19% upon AR8 inoculation (Fig. S4a,b). Collectively, we report a novel PGPF AR8, which demonstrates a robust growth promotion effect on a range of crop species. Thus, the rhizosphere inoculation of AR8 conidia on seedlings of leafy greens was an applicable strategy to maximize the crop growth and yield in such urban crops.

**Fig. 1.**
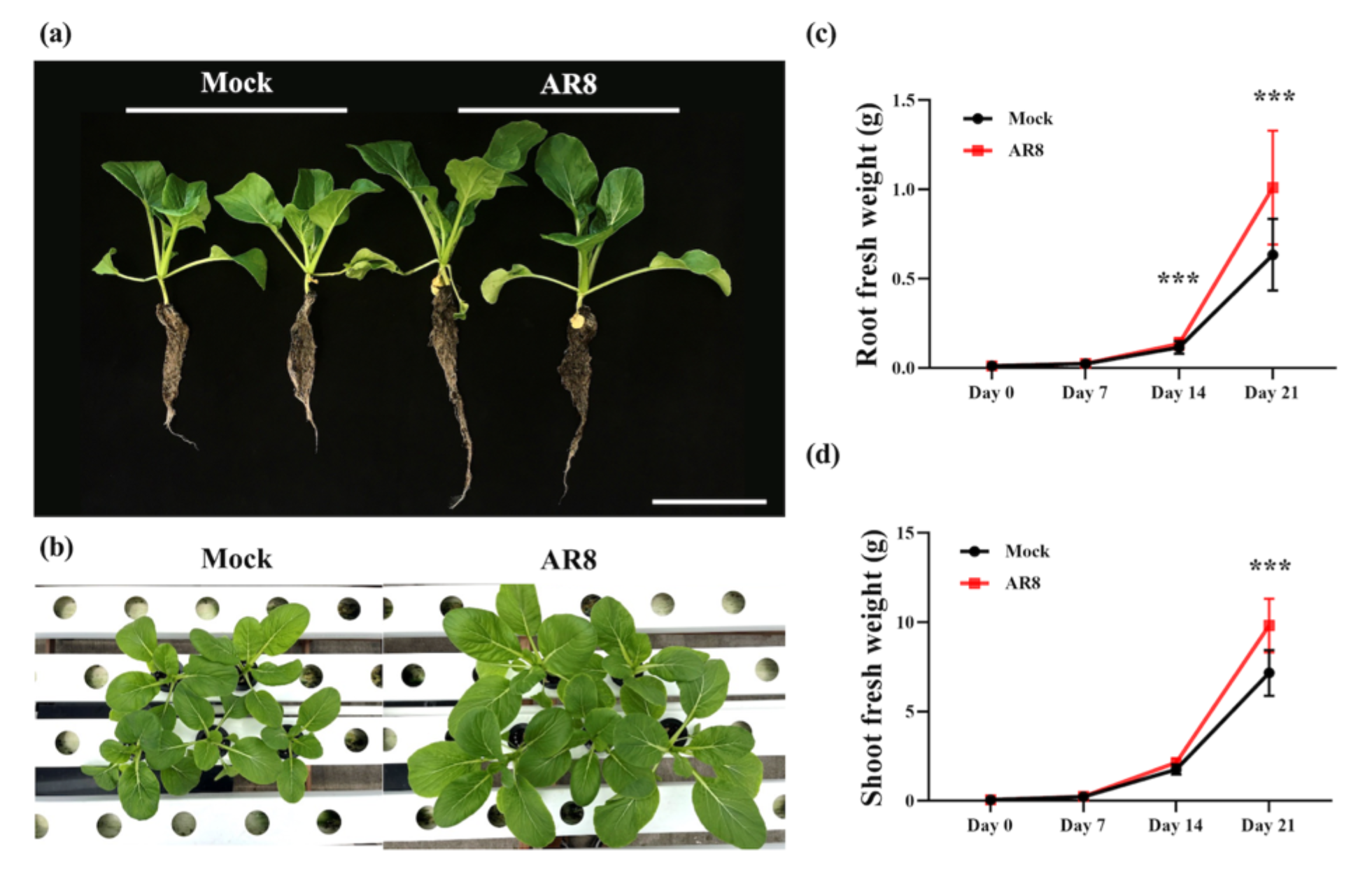
AR8 promotes significant increase in Choy Sum growth under soil conditions. (a) Representative images of Choy Sum seedlings grown in soil with water (mock control) or inoculated with AR8 conidial suspension (10^6^ spores in total) for 21 days. Scale bar, 10 cm. (b-c) Time course (7, 14, and 21 dpi) analysis of Choy Sum shoot (b) and root fresh weight (c) in soil inoculated with water or AR8 conidia (n=12-20 plants per experiment). Data presented (mean ± S.E) was derived from 3 independent replicates of the experiment. Asterisks (***) represent significant differences compared to the mock control at *P* < 0.001 (t-test).

### The nature of AR8 growth and its systemic colonization in Choy Sum roots

To investigate the growth characteristics of AR8, we performed confocal microscopy to first assess the morphology and conidial germination. AR8 conidia produced on PA medium were oblong or short rod-shaped with a uniform length of 4-8 μm (Fig. 2a,b). Interestingly, AR8 conidia germinated and produced apparent germ tubes in water in the presence of Choy Sum roots at 16 and 24 dpi, respectively. By contrast, AR8 conidia failed to germinate in the absence of Choy Sum roots under these conditions (Fig. 2c). This suggested that (Choy Sum) root exudates led to a significant acceleration of conidial germination in AR8, which resembles mycorrhizal symbiosis in being mutually beneficial for both the partners.

**Fig. 2.**
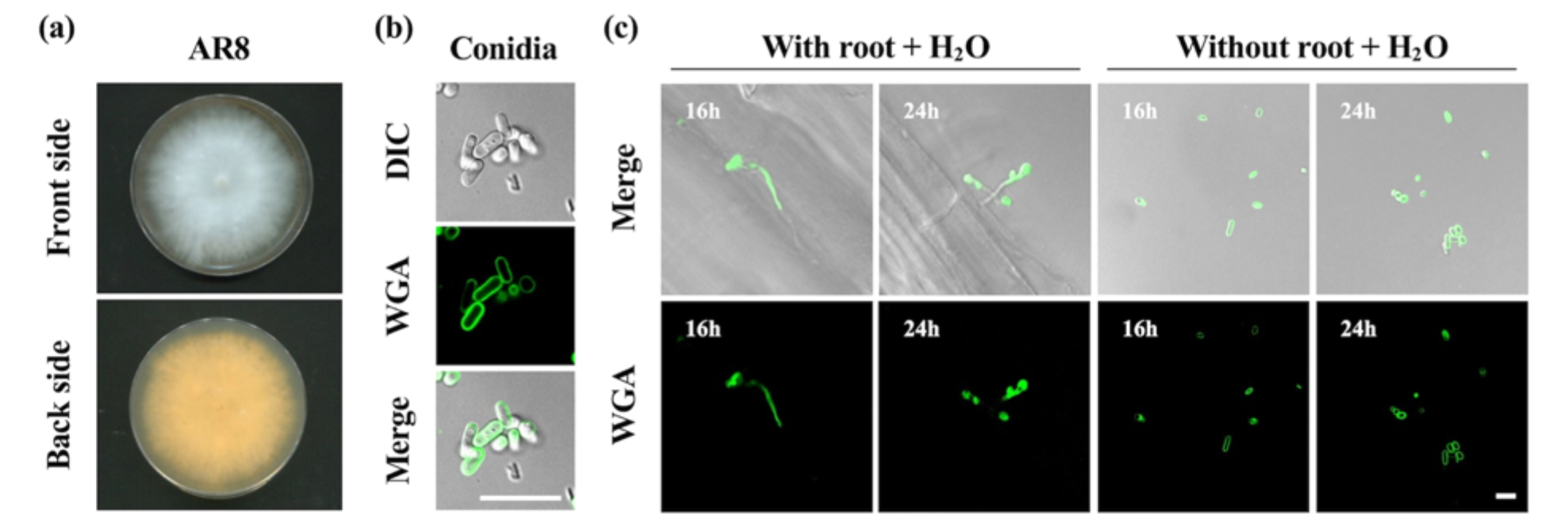
The morphological characteristics of the beneficial fungus AR8. (a-b) AR8 was cultured on PA agar (a) for 5 days and conidia (b) were harvested after subculture under light conditions for 7 days. (c) AR8 conidia were cultured with or without Choy Sum roots for 16 and 24 hours. Laser scanning confocal microscopy of AR8 stained with Wheat Germ Agglutinin-Alexa488 (green; fungal cell wall). Scale bars: 10 μm.

Next, to trace *in planta* colonization process of AR8 by live-cell confocal imaging, we inoculated AR8 conidia on Choy Sum roots and stained with Wheat Germ Agglutinin Alexa Flour^488^ conjugate and Propidium iodide to visualize fungal structures and root cell walls, respectively, under sterilized soil conditions. Tracing the fluorescent signals, indicated a lack of fungal hyphae or mycelial structures in the roots of control samples during *in planta* colonization analysis (Fig. 3a-c). By contrast, Wheat Germ Agglutinin Alexa signal in AR8- inoculated plants indicated that AR8 hyphae attach to the root surface at 4 dpi (Fig. 3d-f). However, at this stage, none of the fungal AR8 hyphae penetrated the epidermal region or colonized the endosphere. Around 7 dpi, AR8 showed a stable interaction with host roots, with fungal hyphae in the rhizosphere gradually enveloping the roots (Fig. 3g). More importantly, AR8 entered the roots interstitially, producing intercellular hyphae that advanced between PI-labeled root epidermal cells (Fig. 3h-j). Thus, we hypothesized that AR8 establishes a biotrophic interaction with plant roots (causing no damage/death, and keeping the host alive) at the early stages of host colonization. Physically intact host root cells clearly outlined by the intercellular hyphae, suggested the viability of host and fungal cells during such intricate AR8 colonization. Following the entry into the root epidermis, AR8 further colonized the root cortex with both inter-and intracellular hyphae at 14 dpi. At this stage, an extensive network of extra-radical hyphae formed and enveloped the host roots (Fig. 3k). The root cortical cells colonized by intracellular hyphae remained unharmed and were inferred to have established a stable biotrophic and symbiotic mycorrhizal interaction within the root system (Fig. 3l,m).

**Fig. 3.**
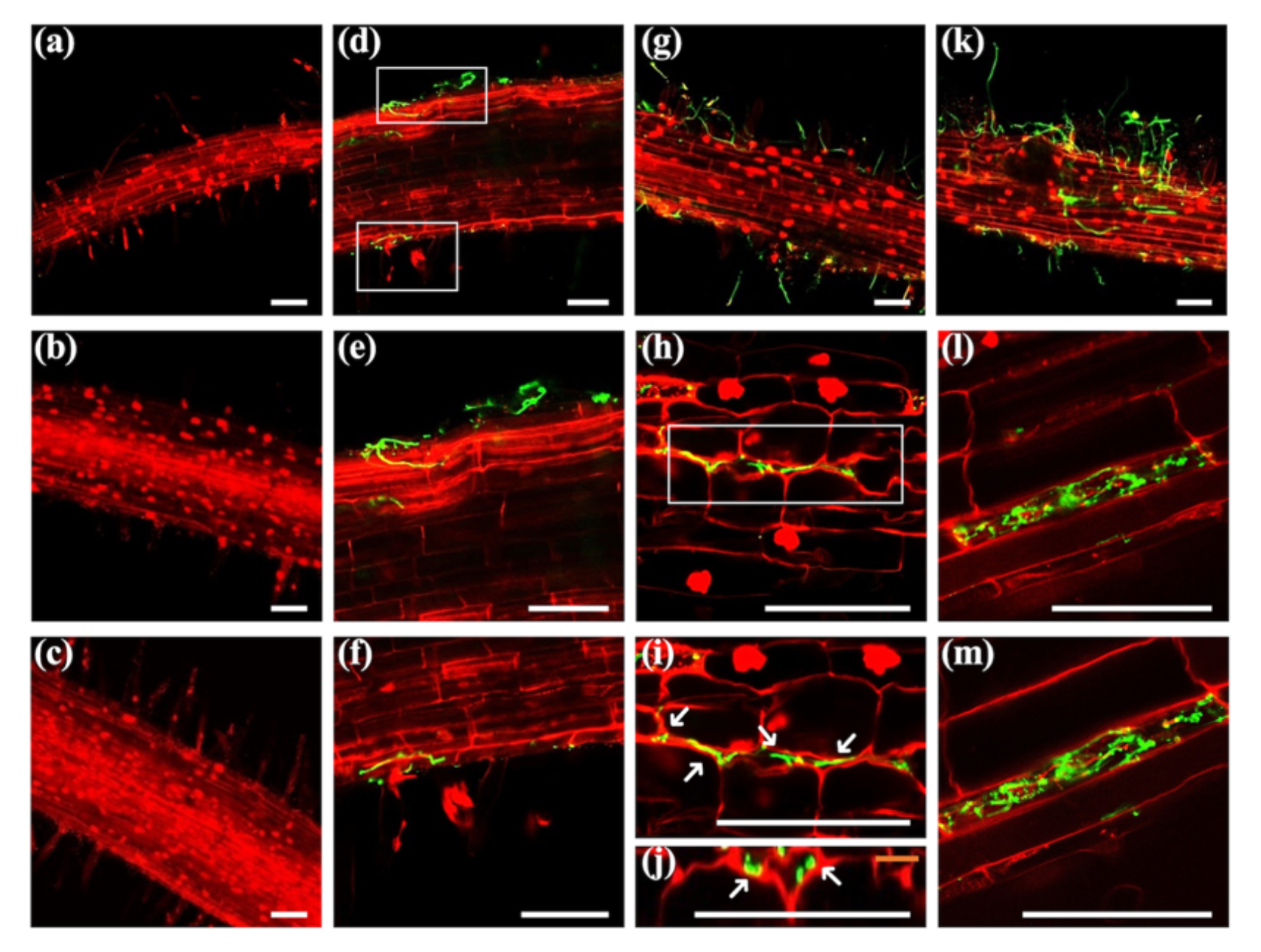
AR8 shows systemic colonization as an endophyte in Choy Sum roots in soil conditions. (a-c) Choy Sum roots were inoculated with water (mock control) and cultivated under sterilized soil conditions for 4 (a), 7 (b), and 14 days (c). No fungal conidia or mycelia are evident around the roots. (d-f) Choy Sum roots inoculated with AR8 conidia (10^6^ spores) in sterilized soil for 4 days. Representative image showing the attachment of AR8 hyphae on the root surface without penetrating the epidermis or inner cortex of the roots (d). Enlarged images of the sections (e-f) from projections shown in (d). (g-j) Choy Sum roots inoculated with AR8 conidia (10^6^ spores) for 7 days. Representative images showing the external (g) and intercellular hyphae (h) of AR8 on the root surface and between root epidermal cells. Enlargement (i) and orthogonal (j) of the sections from the projections is shown in (h). Arrows indicate the intercellular hyphae between Choy Sum root epidermal cells. (k-m) Choy Sum root inoculated with AR8 conidia (10^6^ spores) for 14 days. Representative images of extra-radical hyphae (k) enveloped Choy Sum root and intracellular hyphal (l) colonized in the root cortex. Enlargement of the section (m) from projections shown in (l). Laser scanning confocal micrographs of Choy Sum roots inoculated with water or AR8 conidia and stained with Wheat Germ Agglutinin-Alexa488 and Propidium iodide. Scale bars: 100 μm.

### AR8 promotes plant growth by translocating assimilated phosphorous into the host plants

There have been several studies on fungal endophytes with nutrient solubilizing and/or transport activities, providing evidence that fungus-to-plant nutrient transfer is not only restricted to mycorrhizal symbiosis but is also a common trait between soil fungi and plant associations (Mendes et al., 2013). To address whether AR8 improves plant growth by acquiring and providing soil nutrients, we measured the 3 major macronutrients, nitrogen, phosphorus, and potassium, in Choy Sum under soil conditions by using ICP-MS. We found that AR8 inoculation significantly increased (by about 17%) the phosphorus levels in the shoots whereas the concentration of nitrogen and potassium remained unperturbed therein (Fig. 4).

**Fig. 4.**
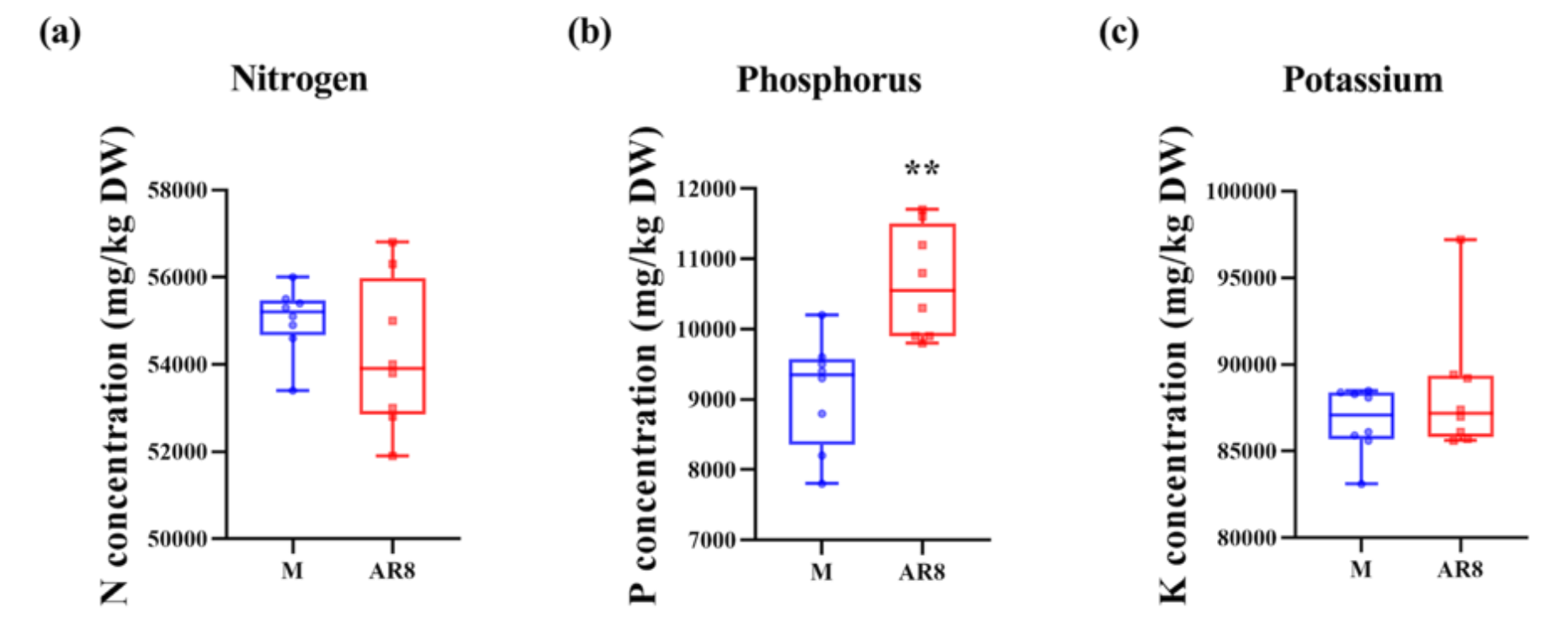
AR8 increases phosphorus accumulation in Choy Sum plants. (a-c) The concentration of nitrogen (a), phosphorus (b), and potassium (c) in shoots of Choy Sum inoculated with water (mock control) or AR8 conidia (10^6^ spores) for 21 days. The nutrient concentration was calculated in mg/kg based on the shoot dry weight (n=16 plants per experiment; three replicates of the experiment). The boxes reveal the first quartile, median and third quartile; the whiskers indicate the minimum and maximum values. Asterisk (**) represents significantly different means compared to the mock control at *P* < 0.01 (t-test).

Fungal hyphae are known to capture environmental Pi outside the rhizosphere and transfer it to the host plants (Bucher, 2007). To verify the Pi acquisition to plant growth in our model, we tested hyphal transport in the bi-compartment system with wild-type and the *phr1* mutant (defective in Phosphate Starvation Response 1) Arabidopsis plants. In this system, AR8 mycelial plugs were inoculated in the inner compartment (HC) while plants were cultivated in the external compartment (RHC) (Hiruma et al., 2016). These two compartments were separated by a plastic barrier, which only allowed the crossing over of AR8 hyphae. By adjusting the Pi concentration in RHC, we aimed to understand whether Pi accumulation in host plants is due to the AR8 hyphal transport. Remarkably, in the low-Pi conditions (100 μM KH_2_PO_4_), the plant size was significantly improved only in the AR8 inoculated wild-type plants. By contrast, the *phr1* mutant defective in Pi starvation response was impaired in such AR8-mediated plant growth promotion (Fig. 5a). Shoot fresh weight of *phr1* mutant plants was comparable between mock and AR8 inoculation (Col-0: 1.52-fold; *P* < 0.001 versus *phr1*: 1.23- fold; *P* = 0.179) (Fig. 5b). Similarly, in the high-Pi conditions (1250 μM KH_2_PO_4_), shoot fresh weight in the AR8-inoculated plants was significantly improved in the wild-type Arabidopsis plants compared to the *phr1* mutant (Col-0: 1.52-fold; *P* < 0.001 versus *phr1*: 1.34-fold; *P* < 0.05), thus indicating the AR8-to-Choy Sum Pi transfer activity (Fig. 5c,d). In summary, we conclude that the transfer of Pi from and/or via AR8 hyphae to the host plants supports shoot growth under both low-and high-Pi conditions, suggesting the functional symbiosis between AR8 and Choy Sum in Pi acquisition and transport.

**Fig. 5.**
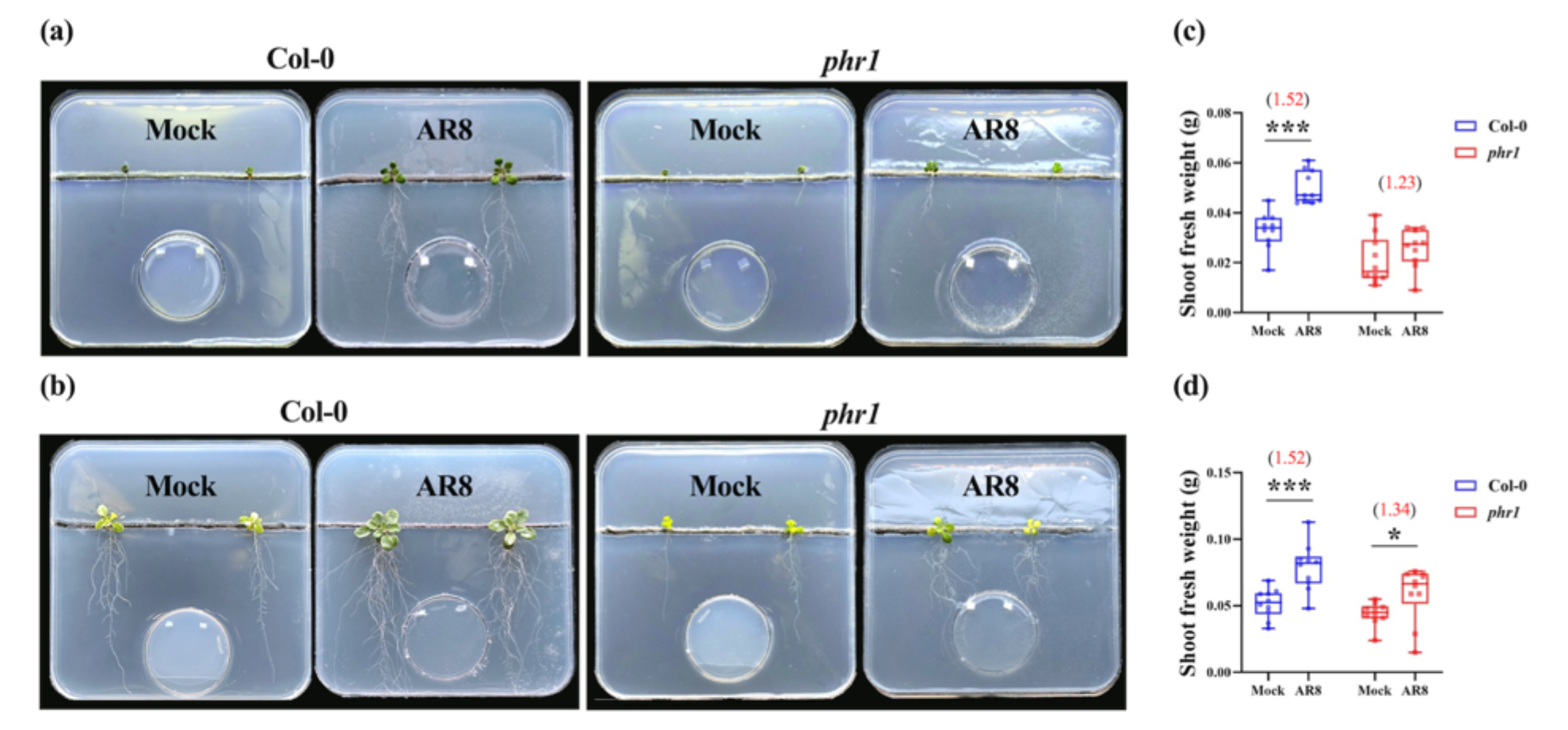
AR8 hyphae are capable of transporting phosphorus to the roots for plant growth promotion. (a-b) Representative images of the bi-compartment system for assessing the Pi transportation. Mock or AR8 mycelial plugs were placed on the MS medium in small round petri dishes (hyphal compartment; HC) of the bi-compartment system while *A. thaliana* Col-0 or the *phr1* mutant seedlings were transferred to MS medium with either low (a) or high Pi (b) in square petri plates (Root hyphal compartment; RHC) of the bi-compartment system. (c-d) Shoot fresh weight of *A. thaliana* Col-0 or the *phr1* mutant line with or without AR8 mycelial plugs in low (c) or high Pi (d) conditions in the bi-compartment system (n=10 plants per experiment; three replicates of the experiment). The boxes reveal the first quartile, median and third quartile, while the whiskers indicate the minimum and maximum values. Asterisks represent significantly different means compared to the corresponding mock control at \**P* < 0.05 and \*\*\**P* < 0.001 (t-test).

To further characterize the Pi solubilizing capacity, AR8 was inoculated in Pikovskaya broth with different inorganic Pi sources (Fig. S5a) (Nautiyal, 1999). AR8 was found to significantly solubilize tricalcium phosphate and hydroxyapatite. The soluble phosphorus concentration was 11.61 ng/ml in tricalcium phosphate and 10.70 ng/ml in hydroxyapatite, respectively (Fig. S5b). Taken together, these results indicate that the formation of extra-radical AR8 hyphal networks likely increases the nutrient absorptive area and rate within the host root system. Furthermore, the Pi solubilizing activity of AR8 was conducive to plant growth and development in natural soils, under low as well as high bioavailable Pi conditions, and thus leading to improved crop productivity.

### AR8 induces global metabolic changes in Choy Sum

To assess the physiological responses by GC-MS-based metabolic profiling (Wagner et al., 2003; Erban et al., 2007), Choy Sum upon mock or AR8 inoculation was performed to identify the changes in primary metabolism in the fungus-inoculated Choy Sum plants in comparison to the mock control. Plants were harvested at the following stages: microgreen stage (7 dpi, first true leaf developed beyond the two cotyledons), seedling stage (14 dpi, with first three true leaves developed after the two cotyledons), and adult stage (21 dpi, in line with harvest time in agricultural practice). Principal component analysis (PCA) and partial least squares-discriminant analysis (PLS-DA) revealed the clustering information between the different groups. Both PCA and PLS-DA showed that the 20 samples (5 biological replicates for each of the 2 treatments at 2 time points) were well-separated and assembled into 4 distinct groups (Fig. S6). The distribution of samples suggested that the shift of metabolic phenotypes was not only across time points but also between treatment regimes.

The time-dependent trajectory analysis demonstrated the metabolic signatures and provided biological insights into plant growth-promoting effect in global (untargeted) metabolomics (Fig. 6a). A total of 309 metabolites were identified based on matches against the NIST mass spectral library. Sugars are products of photosynthesis, and are key metabolites that connect to the tricarboxylic acid (TCA) cycle for energy production (Sheen, 2014). During the vegetative growth phase in Choy Sum, AR8 inoculation strongly increased the levels of most sugars, including monosaccharides (glucose, fructose, and galactose) and sugar alcohols (inositol, xylitol, ribitol, and glycerol). Moreover, TCA intermediates showed higher accumulation in AR8-inoculated Choy Sum, with several metabolites such as citric acid, malic acid, fumaric acid, and succinic acid, showing an increase compared to the mock control.

**Fig. 6.**
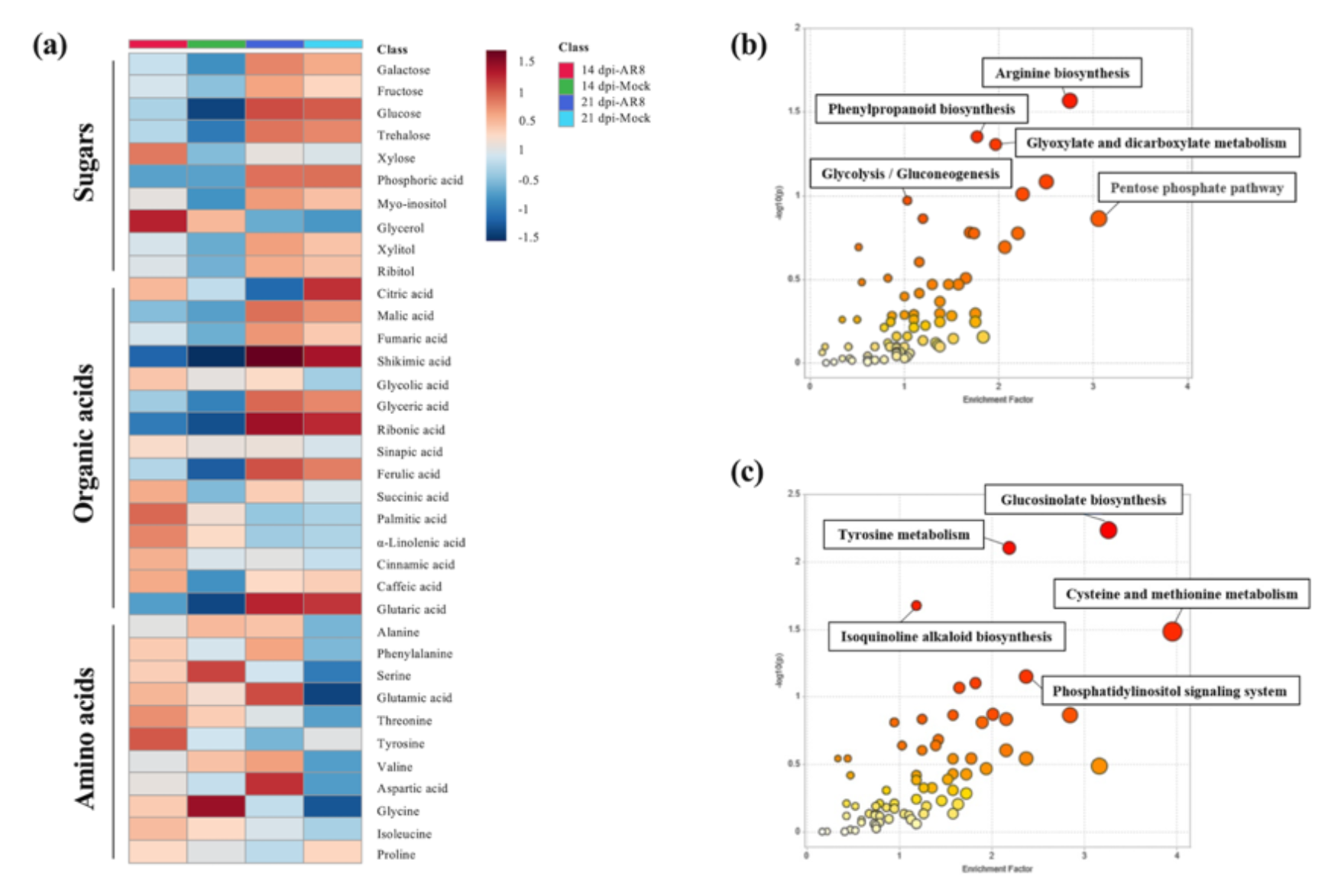
Metabolome profiling and pathway analysis using global (untargeted) metabolomics for Choy Sum. (a) Primary metabolites profiled by GC-EI/TOF-MS and matched to NIST library are displayed as a heatmap for shoots metabolome of Choy Sum at 14 and 21 dpi upon water and AR8 inoculation. The heatmap was generated for Pareto scaling, log transformation, and colored by relative abundance ranging from low (blue) to high (red) via MetaboAnalyst 4.0. Samples (column) are average of five biological replicates while metabolites (rows) are clustered for sugars, organic acids, and amino acids. (b-c) Metabolic pathway analysis in AR8- inoculated plants at 14 (b) and 21 dpi (c) tabled output of metabolic pathway enrichment analysis. The annotated and statistically significant metabolites (*P* < 0.05, fold change > |1.5|) are using for metabolomics pathway analysis. The Y-axis indicates the logP value of the enrichment analysis while X-axis refers to the pathway impact values. The node color is based on the pathway *P* value and the node radius (range 0 to 1, where 1 is maximal impact) displays pathway impact values. Individual nodes represent individual pathways.

Amino acids are primary metabolites with crucial functions as building blocks and/or precursors for the synthesis of nucleic acids, proteins, chlorophyll, and secondary metabolites, regulating key metabolism as well as the formation of vegetative tissues during plant growth and development (Bjorkman et al., 2011; Yang et al., 2020; Trovato et al., 2021). There were significant increases in amino acids levels in Choy Sum upon AR8 inoculation. For example, isoleucine, glycine, and alanine are linked to carbon sources biosynthesis and energy metabolism (Yang et al., 2020). Increased accumulation of these 3 amino acids was observed at both the seedling and adult stages of AR8-inoculated Choy Sum plants.

In the detailed (targeted) analysis, eight sugars, including four monosaccharides (glucose, fructose, mannose, and galactose), a disaccharide (sucrose), and three sugar alcohols (inositol, erythritol, and mannitol) were measured (Fig. S7a). Carbohydrate metabolism was significantly impacted, and in general, we further confirmed the increased accumulation of these primary metabolites in AR8-inoculated Choy Sum. Accumulation of glucose and fructose, the two major monosaccharide carbon sources, was significantly increased during vegetative growth (Fig. 7a,b), while other monosaccharides and sugar alcohols were also observed with significant changes (Fig. S7a). By contrast, the disaccharide, sucrose, did not show significant changes during vegetative growth of AR8-inoculated and control plants (Fig. 7c). On the other hand, the concentration of 19 proteinogenic amino acids were quantified, for which cysteine was found to be lower than the detection limit, and hence was excluded from the analysis (Fig. S7b). AR8 inoculation led to an increased accumulation of amino acids directly related to energy-producing metabolism (Hildebrandt et al., 2015; Yang et al., 2020). These include lysine, isoleucine, methionine, and alanine, which showed significant accumulation at microgreen and adult stages of growth in the host plants (Fig. 7d-g). Besides, aromatic amino acids (phenylalanine, tyrosine, and tryptophan) in the AR8-inoculated Choy Sum showed significant differences during vegetative growth, except for tryptophan in microgreens and seedlings and tyrosine in seedlings only (Fig. S7b). Aromatic amino acids are essential components for the synthesis of secondary metabolites with multiple biological functions (Tzin and Galili, 2010). This observed accumulation in Choy Sum indicates an increased rate/induction of secondary metabolism pathway(s) upon interaction with AR8. In summary, our results reveal the profile of various primary metabolites at different growth stages of Choy Sum, and further provide a metabolic link between AR8 and the associated plant growth promotion effect. The increase in the levels of glucose and fructose suggests that the increased pool of carbon sources is likely an attempt to promote core metabolism for energy production. Higher concentrations of sugar alcohols and growth-related amino acids also reflect the scenario of greater energy-producing and growth-promoting metabolism enabled by *Tinctoporellus* AR8 strain in Choy Sum plants.

**Fig. 7.**
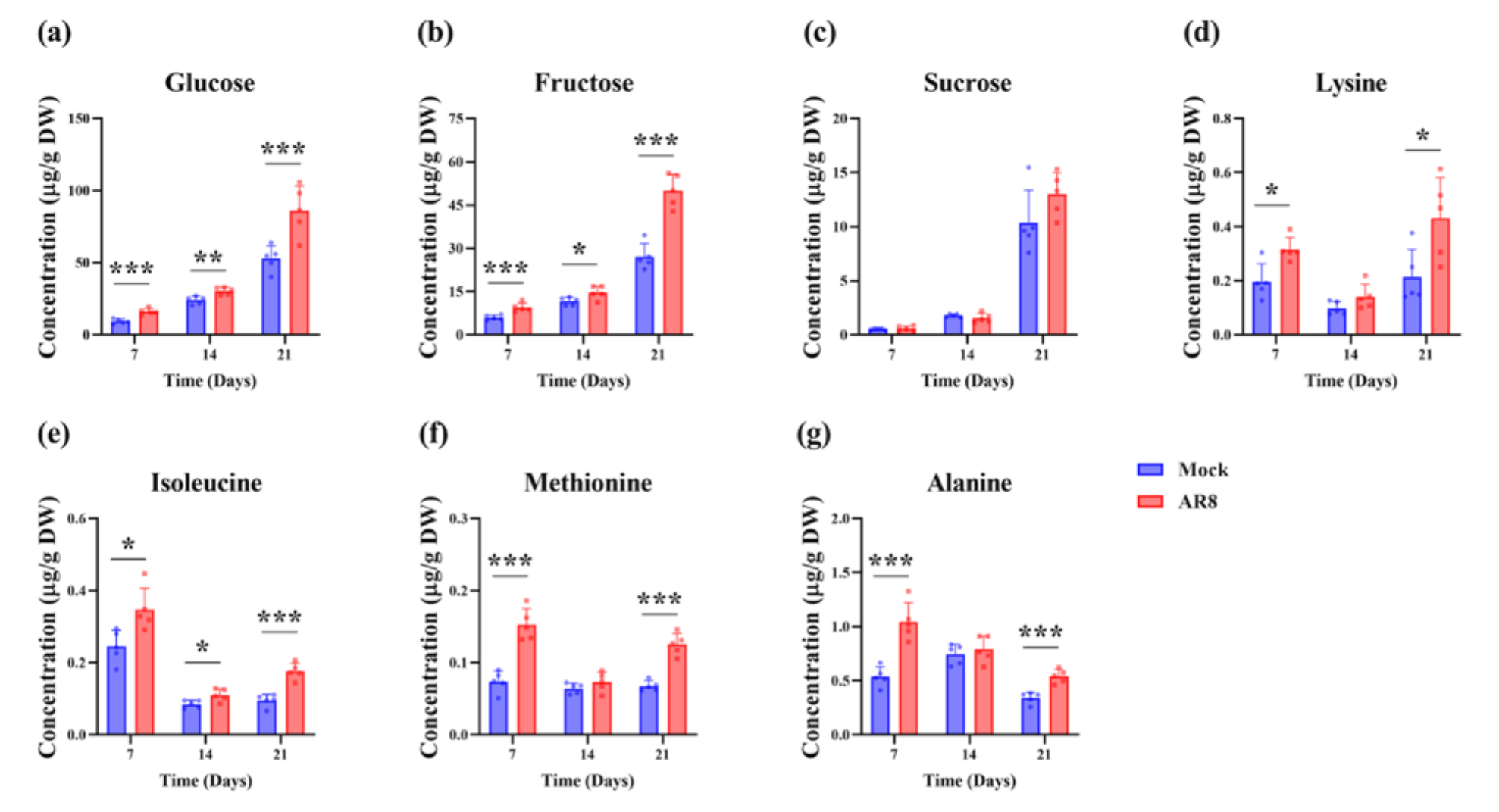
AR8 increases the accumulation of sugars and amino acids during vegetative growth of Choy Sum. (a-c) Quantification of glucose (a), fructose (b), and sucrose (c) in in shoots of Choy Sum inoculated with water (mock control) or AR8 conidia (10^6^ spores in total) for 7 (microgreen), 14 (seedling), and 21 dpi (adult) grown in soil. (d-g) Quantification of lysine (d), isoleucine (e), methionine (f), and alanine (g) in in shoots of Choy Sum inoculated with water or AR8 conidia for 7, 14 and 21 dpi grown in soil. Choy Sum shoots (n=16 plants per experiment) of each group were pooled to measure metabolite concentration. The concentration was calculated in μg/g based on the shoot dry weight. Data presented (mean ± S.E) were derived from 3 independent replicates of the experiment. Asterisks represent significant differences compared to AR8 and corresponding mock control at \**P* < 0.05, \*\**P* < 0.01, \*\*\**P* < 0.001 (t-test).

### AR8 induces the phenylpropanoid *t*-CA as a growth-promoting metabolite in Choy Sum

The mode of action of a fungal endophyte on plant growth is complex and unable to be accurately described in terms of just a set of metabolites. Instead, a wide range of direct or indirect metabolic targets interacting with multiple enzymes or pathways characterizes the response of PGPF in plant growth and development. Thus, we further evaluated the metabolic changes of AR8 to plant growth promotion by pathway analysis. The annotated and statistically significant (*P* < 0.05, fold change > |1.5|) metabolites detected in the untargeted metabolomics analysis were subjected to such pathway classification/predictions using MetaboAnalyst 4.0 (https://www.metaboanalyst.ca/MetaboAnalyst/home.xhtml) (Xia and Wishart, 2011). The results showed that the metabolic pathways most likely to be reprogrammed upon AR8 inoculation in Choy Sum were: arginine biosynthesis, phenylpropanoid biosynthesis, glyoxylate and dicarboxylate metabolism, glycolysis/gluconeogenesis, and pentose phosphate pathway at 14 dpi (Fig. 6b); and glucosinolate biosynthesis, tyrosine metabolism, isoquinoline alkaloid biosynthesis, cysteine and methionine metabolism, and phosphatidylinositol signaling systems at 21 dpi (Fig 6c). Characterized below are the significant changes evident in phenylpropanoid biosynthesis during *Tinctoporellus*-Choy Sum interaction (Fig. S8). We focused on this pathway since it is critical to plant-environment interactions, i.e., metabolites of phenylpropanoid pathway contribute to the priming of plant defense (Dixon et al., 2002; Bressan et al., 2009). Moreover, recent findings have provided a potential link between phenylpropanoid biogenesis and plant growth (Brown et al., 2001; Kurepa et al., 2018). Therefore, observed changes at the seedling stage of Choy Sum upon AR8 inoculation prompted us to explore the role of specific phenylpropanoids in such beneficial plant-fungus interactions.

Phenylalanine derived from shikimic acid is the starting point for the phenylpropanoid pathway, which can be converted into *t*-CA and diverted towards the biosynthesis of a number of polyphenol classes, including lignin and flavonoids. These are reported to function in several plant growth and developmental processes and are metabolic signatures in mycorrhizal associations (Kaur and Suseela, 2020; Dong and Lin, 2021). In AR8-plant interaction, Shikimic acid was significantly increased at the microgreen and seedling stages in Choy Sum (Fig. S8). Hydroxycinnamic acids (*t*-CA, caffeic acid, and ferulic acid) showed a different trend in contrast as they showed a significant increase in the seedlings or/and adult Choy Sum plants, albeit with a dramatically decreased accumulation at the microgreen stage (Fig. S8). Lignin is derived from hydroxycinnamic acid and is one of the major components of the plant cell wall.

As the main pathway involved in plant cell wall composition or strengthening, lignin plays a vital role in plant growth, defense response, and environmental adaptation (Dong and Lin, 2021). Intermediates of lignin biosynthesis in which caffeic acid and ferulic acids serve as precursors, including coniferaldehyde, sinapic acid, caffeoylquinic acids, and feruloylquinic acids also significantly increased in Choy Sum during vegetative growth in the presence of AR8 (Fig. S8). In summary, quantification of hydroxycinnamic acids and their derivatives confirmed the patterns observed in the untargeted metabolomics and pathway analyses, providing further evidence for the reprogramming of phenylpropanoid biosynthesis upon and during prolonged association of AR8 with the host plants.

Intriguingly, the increase of hydroxycinnamic acids at the seedling stage of Choy Sum upon AR8 inoculation was concomitant with the observed dramatic growth promotion effect. Hence, we hypothesized that hydroxycinnamic acids triggered by AR8 contribute to induction of plant growth. To characterize the impact of hydroxycinnamic acids on plant growth, Choy Sum seedlings were grown in MS medium supplemented with either *t*-CA, *p*-coumaric acid, caffeic acid, or ferulic acid (Fig. S9). As expected, seedlings grown in the medium supplemented with *t*-CA showed a growth-promoting effect on shoot biomass (an increase of 28.30%) while seedlings grown in other hydroxycinnamic acids did not. Subsequently, the growth-promoting activity of *t*-CA was tested in a dose-dependent manner on Choy Sum seedlings (Fig. S10). A clear trend towards growth promotion was only observed in shoot biomass of Choy Sum (an increase of 22.93%, 38.01%, and 24.69%, respectively) upon treatment with low concentration (1, 2.5, and 5 μM) of *t*-CA. By contrast, the growth inhibitory effect of exogenous *t*-CA at high concentration was evident in Choy Sum whereby a strong decrease in shoot biomass (18.42% and 76.16%, respectively) was observed upon treatment with 10 and 50 μM *t*-CA.

To gain insight into the growth-promoting effect of *t*-CA in the AR8 model in Choy Sum, seedlings were cultivated in the presence of a low dose (1 or 2.5 μM) of *t*-CA and AR8 separately and/or in combination (Fig. 8a,b). Importantly, a further enhancement was observed in Choy Sum when *t*-CA and AR8 were applied together. Compared with AR8 inoculation alone, the combined treatment (1 μM *t*-CA+AR8 and 2.5 μM *t*-CA +AR8) led to a further growth increase of 17.99% and 47.69% on Choy Sum, respectively. These results suggest an additive effect or positive interaction between *t*-CA-dependent and AR8-mediated plant growth-promoting mechanisms.

**Fig. 8.**
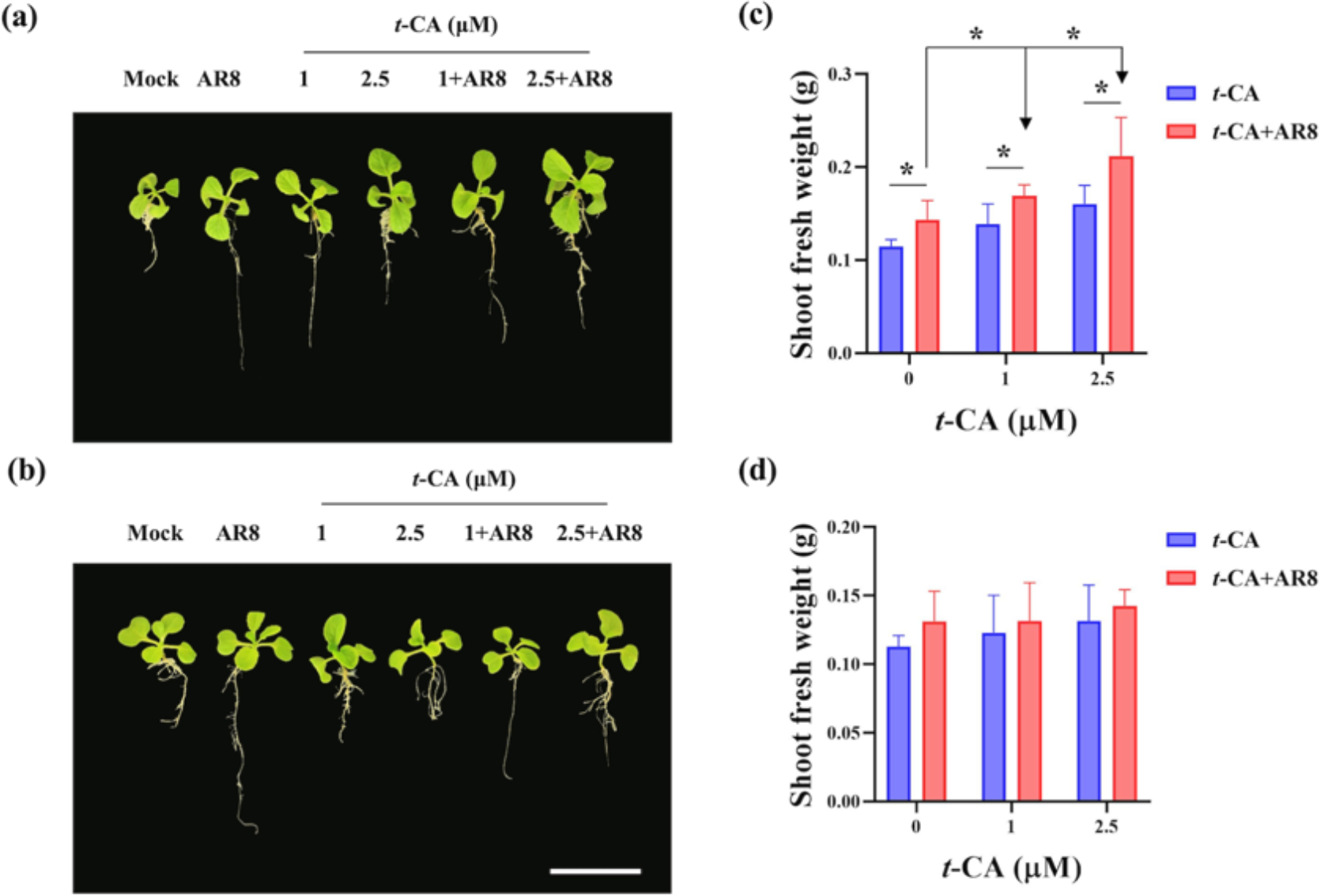
Light source and quality affects growth promotion effect of *t*-CA and AR8 in Choy Sum seedlings. (a-b) Representative images of *t*-CA application in low concentration and/or in combination with AR8 on Choy Sum. Choy Sum seedlings were grown in MS medium inoculated with either *t*-CA (1 and 2.5 μM), AR8 mycelial plugs, or in combination under tube light (a) or LED lighting conditions (b) for 7 days. Scale bar, 5 cm. (c-d) Shoot fresh weight of Choy Sum seedlings (n=5 plants per experiment; three replicates of the experiment) upon AR8 inoculation, *t*-CA treatment (1 and 2.5 μM), or denoted concentration of *t*-CA with AR8 in combination under tube light (c) and LED lighting conditions (d). Data presented (mean ± S.E) were derived from 3 independent replicates of the experiment. Asterisks represent significant differences compared to AR8 and corresponding mock control at \**P* < 0.05 (t-test).

CA exists as a *cis* or *trans* isomer in plants. Notably, the isomerization of CA is driven by UV-B light. The duration and intensity of light directly determines the efficiency of such isomer conversion. In previous findings, UV-B from the incandescent tube lights could establish a 60/40 *cis*/*trans*-CA equilibrium throughout the experiment in plant growth chambers (Steenackers et al., 2017). To explore whether isomerization of CA contributes to growth promotion in plant-fungus interactions, Choy Sum seedlings were cultivated as described above under LED light conditions (Fig. 8c,d). In the absence of UV-B in the LED spectrum, seedlings grown in the medium in the presence of *t*-CA and AR8 separately and/or in combination did not show any growth induction or increase in shoot biomass. These results helped us reach two conclusions. First, *c*-CA is likely the bioactive compound providing plant growth-promoting activity. Second, since shoot biomass is greatly reduced under LED conditions, it is more likely that AR8 increases and utilizes phenylpropanoid metabolism including cinnamic acid biosynthesis to stimulate plant growth.

### AR8 is a PGPF that confers beneficial effects to Choy Sum: from macro-to micro-scale

In summary, we propose a model integrating the beneficial mechanisms of AR8, a new fungal endophyte identified from the rhizosphere of Arabidopsis. Typically, the colonization by AR8 initiates on the root surface but it can also systemically colonize both intra-and intercellular regions of Choy Sum roots. These extra-radical hyphae and the Pi solubilizing activity of AR8 enhance soil exploration capacity and promote higher plant nutrition in Choy Sum. Proceeding to the molecular mechanism(s) of plant growth promotion, the reprogramming in the shoot metabolome further ensures the beneficial phenotype following the plant-fungal interaction. The higher abundance of sugars, amino acids, and phenylpropanoids reveals the metabolic signatures in Choy Sum upon AR8 inoculation, which resembles functional mycorrhizal symbiosis. More importantly, we used the metabolic signatures to identify a specific metabolite and characterized its plant growth-promoting function. Our results support a model explaining the increase in shoot biomass as the consequence of higher *t*-CA levels upon AR8 inoculation. Our study also highlights the beneficial activity of AR8 on different crop species. This emphasizes the necessity to further explore the host range and plant growth-promoting mechanism(s) (e.g., phytohormones) in plant-AR8 beneficial mycorrhizal associations for improved crop production.

## Discussion

In this study, AR8 was reported to be a PGPF in the urban farm crop Choy Sum (Figs 1, S11). The phylogenetic analysis of AR8 revealed a close relationship to other *Tinctoporellus* species in particular *epimiltinus* based on the ITS-based rDNA sequence analysis. However, little information is currently available on the colony characteristics of *Tinctoporellus* and/or its phylogeny. It was first circumscribed as a monospecific genus by Ryvarden and *T. epimiltinus* was described as a type species (Ryvarden, 1979). Thereafter, *T. isabellinus*, *T. bubalinus*, and *T. hinnuleus* were added to this genus in 2003 and 2012, respectively (Ryvarden and Iturriaga, 2003, Yuan and Wan, 2012). To gain more insights into the biological functions of AR8-plant interaction, further studies will focus on the whole-genome sequencing, annotation and analysis of AR8 genome and the identification of transcriptome in the host plants to elucidate the molecular correlation with the plant growth-promoting metabolism reported here for this beneficial mycorrhizal endophyte.

In the AR8-Choy Sum interaction, Choy Sum provides the required nutrient niche to improve the germination of AR8 conidia and promote the establishment of such beneficial interaction in the rhizosphere (Fig. 2). Intriguingly, the root colonization by AR8 appeared to be a more prolonged process compared to other fungal endophytes. Taking *C. tofieldiae* as an example, it only takes 2 days for *C. tofieldiae* to initiate biotrophic interaction with inter-and intracellular hyphae in the root endosphere of *A. thaliana* (Hiruma et al., 2016). In comparison, from hyphal attachment to the intracellular endophytic growth pattern, AR8 took around 1-2 weeks to establish a stable interaction within the Choy Sum roots (Fig. 3). Thus, we predict that the beneficial impact of AR8 on the host plant growth already initiates/occurs during the early stages of colonization. The molecular and biochemical mechanisms in host plants are likely to be highly activated or induced and continuously improved plant growth and development from hyphal attachment that precedes the initial intercellular hyphal growth or interstitial interaction around the cell-cell junctions in the roots. If we refer to the growth promotion assay (Figs 1b, S2a), there is no significant increase in shoot biomass until the formation of intracellular AR8 hyphae in Choy Sum roots at 7-14 dpi (Fig. 3l, 3m). These results partially confirmed our predictions and helped us propose the spatiotemporal dimension of the endophytic colonization process associated with the plant growth promotion effects by AR8. Overall, we suggest that AR8 is a beneficial mycorrhizal endophyte, which colonizes the intra-and intercellular compartments of the host roots and provides significant beneficial effects to the plants without causing any apparent damage or detrimental symptoms.

Plant growth promotion by AR8 was shown to be associated with Pi acquisition. AR8 was able to transport Pi to plants under high-and low-Pi conditions (1250 and 100 μM KH_2_PO_4_, respectively) on MS agar medium (Fig. 5). This stands in stark contrast to the fungal endophyte *C. tofieldiae*, in which beneficial activities are restricted strictly to the low-Pi conditions (50 μM KH_2_PO_4_) (Hiruma et al., 2016). In accordance with the role of AR8 in Pi acquisition, the increased nutrient absorption from the soil implies the potential use of AR8 as a biofertilizer in agriculture. Indeed, the colonization of AR8 reflects an expanded capacity of host roots to access Pi by the long-distance transport of/via fungal hyphae. In addition to improving soil exploration capacity through fungal hyphae, the solubilization of plant-inaccessible Pi might be one of the relevant mechanisms by which AR8 accesses environmental P sources. Notably, an *in vitro* study of AR8 indicates that the beneficial activity of AR8 involves the solubilization of inorganic Pi (tricalcium phosphate and hydroxyapatite) sources (Fig. S5). Although the insoluble Pi solubilization activity of AR8 was lower than that of the known PGPF B9 (Gu et al., 2023), it convincingly supports the notion that AR8 expands the range of nutrient absorption by Pi solubilization and hyphal transport, thereby promoting plant growth. Taken together, our research provides evidence that fungus-to-plant Pi transport, generally attributed to mycorrhizal interactions, is also found in the AR8 model.

The metabolic profiling helped gain molecular insights into the AR8-Choy Sum interaction. A marked increase in the levels of primary metabolites suggests an improved plant growth performance upon AR8 inoculation (Figs 6a, 7, S7). Sugars are the key components that reflect energy status and, therefore, the capacity to respond to sugar levels is critical for plants to maintain an appropriate physiological state (Lastdrager et al., 2014). Further increases in the content of various sugars in AR8-inoculated Choy Sum during vegetative growth provide evidence for the advanced photosynthetic capacity and the idea of plant growth promotion (Figs 7a-c, S7a). On the other hand, in plant-mycorrhizal symbiosis, host plants reciprocate the nutrient supply of AMF by providing sugars and fatty acids (Schweiger and Müller, 2015; Jiang et al., 2017). Thus, higher concentrations of sugars found in the leaves of mycorrhizal plants suggest that AMF reprogram the sugar metabolism as an upgrade of carbon source and regulate the sugar accumulation in roots to meet their demands (Lendenmann et al., 2011). In addition to plant metabolome, trehalose is a fungus-specific sugar converted from hexose (e.g., glucose and fructose) assimilated by AMF in the intra-radical hyphae, which can be linked to a functional mycorrhizal symbiosis (Bago et al., 2000; Lohse et al., 2005). In the AR8 model, the higher relative abundance of trehalose in AR8-inoculated Choy Sum in global metabolomics indicates a potential carbon drain from the host plants to AR8 (Fig. 6a).

Apart from sugars, AR8 also impacts the TCA cycle and amino acid biosynthesis in Choy Sum (Figs 6a, 7d-g, S7b). The activation of the TCA cycle provides the adenosine triphosphate and carbon skeletons necessary for amino acid production (Lohse et al., 2005). These increased metabolites help plants in their growth as well as adaptation to the environment. Among the mentioned compounds, *t*-CA was brought to the front as an essential factor in the AR8 model. In plant-mycorrhizal interactions, a higher abundance of hydroxycinnamic acids and their derivatives is recognized as the priming of the defense system of the plant. On the other hand, as defense molecules, a massive accumulation of these metabolites may negatively affect mycorrhizal colonization due to their antimicrobial properties (Maier et al., 2000; Aliferis et al., 2015). Thus, in our case, the plant growth-promoting activity of cinnamic acid during the experiment is likely to be unintentional. The accumulation of *t*-CA at the seedling stage of AR8- inoculated Choy Sum was likely exposed to environmental UV-B, thereby converting to *c*-CA and promoting plant growth (Fig. 8). Cinnamic acid is claimed to be a growth-promoting compound in *A. thaliana* (Kurepa et al., 2018). The narrow dose range in which cinnamic acid acts as a growth stimulant may be a general effect not only in Choy Sum but also in other plant species (Fig. S10). *c*-CA is the most promising candidate that can alter auxin transport and affect plant growth among metabolites in phenylpropanoid pathways (Steenackers et al., 2019). However, whether and at what level auxin signaling interacts with *c*-CA for plant growth promotion in plant-fungal interaction remains to be tested. The possibility of *c*-CA being involved in auxin-independent mechanisms cannot be excluded from the AR8 model, although auxin seems to have a crucial role based on previous findings. Regardless of the hypothesis, the fact that phenylpropanoid cinnamic acid induction by AR8 governs the developmental control of Choy Sum growth, suggests an intricate and multifaceted plant-mycorrhiza interaction that influences overall growth and fitness in the host.

Taken together, despite fungi not being the major microbiome constituents in the rhizosphere, we cannot overlook their contribution to plant growth and health. In our study, AR8 demonstrated a biotrophic lifestyle and colonized the root inter-and intracellular compartments without causing negative influences and led to increased plant biomass. More importantly, the Choy Sum-AR8 interaction provides an excellent example of a symbiotic partnership for exploring such poorly understood mycorrhizal associations. Lastly, increased molecular understanding of the biology of host-mycorrhizal endophyte systems will further the adoption and application of such biofertilizers in sustainable-and precision agriculture.

## Acknowledgements

We thank Poonguzhali Selvaraj, Wenhui Zheng, and Keyu Gu for their help at the initial stages of this project. We thank the Fungal Patho-Biology Group for discussions and suggestions. We are grateful to the Chua Lab (Chung-Hau Hwang) for sharing the Arabidopsis *phr1* mutant line; and to Yang Fan for technical support in confocal imaging. Our sincere thanks to the Sanjay Swarup Lab, and the NUS Environmental Research Institute (Singapore) for help in metabolomics analysis.

## Author Contributions

C.-Y.C., and N.I.N. designed the experiments. C.-Y.C., performed all the experiments. C.-Y.C., and N.I.N. interpreted and analyzed the data, compiled all the results, and co-wrote the manuscript. N.I.N. provided funding support and resources, and overall project management. All authors agree to the final submitted version of the manuscript.

---

**Fig. S1** Fungal isolates from Arabidopsis rhizosphere modulate Choy Sum growth. (a-b) Shoot fresh weight of Choy Sum plants inoculated with non-PGPF isolates (AR1, AR2, AR4, AR12, AR13, AR14, AR19, AR22, AR30, AR36, AR50, and AR51) (a) or PGPF isolates (AR8, AR9, AR11, AR18, AR32, AR36, AR51, AR65, and AR70) (b) in soil conditions. Choy Sum seedlings (n=8 plants per experiment; three replicates of the experiment) inoculated with water (mock control) or conidia (10^6^ spores in total) from the indicated fungal isolate in each instance, and shoot fresh weight measured at 21 days post-inoculation (dpi). M refers to mock control inoculated with water. Numbers in (b) represent the fold change in shoot fresh weight between fungal inoculation and mock control. The boxes reveal the first quartile, median and third quartile; the whiskers indicate the minimum and maximum values. Asterisks represent significantly different means compared to the mock control at \**P* < 0.05, \*\**P* < 0.01, \*\*\**P* < 0.001 (Student’s t-test).

**Fig. S2** AR8 promotes plant growth and fertility under soil conditions. (a) Representative images of Choy Sum seedlings grown in soil with water (mock control) or AR8 conidia (10^6^ spores) for 7 and 14 days. Scale bar, 10 cm. (b) Representative images of floral initiation time point in mock control or AR8-inoculated Choy Sum plants. Images shown are at 28 dpi. (c) Silique number produced by Choy Sum grown in soil measured at 49 dpi. Data presented (mean ± S.E) was derived from 3 independent replicates of the experiment. The boxes reveal the first quartile, median and third quartile; the whiskers indicate the minimum and maximum values. Asterisks (*) represent significantly different means compared to the mock control at *P* < 0.05 (t-test).

**Fig. S3** AR8 promotes growth in Brassicaceae plants under soil conditions. (a-b) Representative images of *Arabidopsis thaliana* Col-0 (a) and Kailan (b) grown in soil with water (mock control) or AR8 conidia (10^6^ spores) for 21 days. Scale bars, 10 cm. (c-d) Shoot fresh weight of *Arabidopsis thaliana* Col-0 (c) and Kailan (d) grown in soil inoculated with water or AR8 conidia (n=12 and 24 plants per experiment, respectively; three replicates of the experiment). The boxes reveal the first quartile, median and third quartile; the whiskers indicate the minimum and maximum values. Asterisks represent significantly different means compared to the mock control at \*\**P* < 0.01, \*\*\**P* < 0.001 (t-test).

**Fig. S4** AR8 promotes growth in cereal crops under soil conditions. (a-b) Representative images of rice cultivar CO39 (a) and barley cultivar Express (b) grown in soil with water (mock control) or AR8 conidia (10^6^ spores) for 21 days. (c-d) Shoot fresh weight of rice cultivar CO39 (c) and barley cultivar Express (d) grown in soil inoculated with water or AR8 conidia (n=20 plants per experiment, respectively; three replicates of the experiment). The boxes reveal the first quartile, median and third quartile; the whiskers indicate the minimum and maximum values. Asterisks represent significantly different means compared to the mock control at \**P* < 0.05 (t-test).

**Fig. S5** AR8 solubilizes inorganic Pi sources in Pikovskaya broth. (a) Pi solubilizing capability of AR8 and B9 in different inorganic Pi sources (tricalcium phosphate, hydroxyapatite, aluminum phosphate, and iron phosphate) after 7 days. (b) Soluble Pi concentration produced by AR8 and B9 in in Pikovskaya broth with different inorganic Pi sources. Mycelial plugs (AR8 and B9) were inoculated to Pikovskaya broth with different inorganic Pi sources for 7 days and the soluble Pi concentration was measured using phosphomolybdenum spectrophotometry. Data presented (mean ± S.E) was derived from 3 independent replicates of the experiment. Asterisks represent significant differences compared to the mock control at \*\*\**P* < 0.001 (t-test).

**Fig. S6** Overview of metabolic changes in Choy Sum upon AR8 inoculation. (a-b) The PCA (a) and PLS-DA (b) score plots for shoots metabolome of Choy Sum plants inoculated with water or AR8 conidia (10^6^ spores) at 14 and 21 dpi as captured by GC-EI/TOF-MS. The colored ellipses represent 95% confidence regions for each group.

**Fig. S7** Repertoire of primary metabolites (sugars and amino acids) accumulating in Choy Sum during beneficial association with AR8. (a-b) Heatmap showing quantitative and qualitative changes in concentration of sugars (a) and amino acids (b) in Choy Sum shoots at three growth stages (microgreen, seedling, and adult) upon water or AR8 conidial inoculation were performed by targeted analysis. The heatmap was generated through log transformation and colored by concentration (μg/g), e.g., 10 becomes 1, highlighted in blue (low) or red (high) via GraphPad Prism 8, respectively. Numbers represent the average concentration of corresponding metabolites derived from five biological replicates.

**Fig. S8** Repertoire of phenylpropanoid (hydroxycinnamic acids) accumulating in Choy Sum during beneficial association with AR8. Heatmap showing quantitative and qualitative changes in abundance of hydroxycinnamic acids and their derivatives in Choy Sum shoot at three growth stages (microgreen, seedling, and adult) upon water or AR8 inoculation were performed by targeted analysis. The heatmap was generated through log transformation and colored by concentration (μg/g), e.g., 10 becomes 1, highlighted in blue (low) or red (high) via GraphPad Prism 8, respectively. Numbers represent the average concentration of corresponding metabolites abundance of five biological replicates.

**Fig. S9** Impact of specific hydroxycinnamic acids on the growth of Choy Sum. (a) Representative images of Choy Sum seedlings grown in MS medium containing *t*-CA (2.5 μM), *p*-coumaric acid (60 μM), ferulic acid (20 μM), and caffeic acid (20 μM) for 7 days. Scale bar, 5 cm. (b) Shoot fresh weight of Choy Sum grown on control and hydroxycinnamic acids-supplemented (*t*-CA, *p*-coumaric acid, ferulic acid, and caffeic acid) medium for 7 days (n=10 plants per experiment). Data presented (mean ± S.E) were derived from 3 independent replicates of the experiment. Asterisks represent significant differences compared to the mock control at \*\**P* < 0.01 (t-test).

**Fig. S10** *trans-*Cinnamic acid displays growth-promoting activity in Choy Sum plants in a dosage-dependent manner. (a) Representative images of Choy Sum seedlings grown in MS medium containing the denoted amounts of *t*-CA (0, 1, 2.5, 5. 10, and 50 μM) for 7 days. Scale bar, 5 cm. (b) Shoot fresh weight of Choy Sum grown on uninoculated control or *t-*CA-supplemented medium for 7 days (n=10 plants per experiment). Data presented (mean ± S.E) were derived from 3 independent replicates of the experiment. Asterisks represent significantly different means compared to the mock control at \**P* < 0.05, \*\**P* < 0.01, \*\*\**P* < 0.001 (t-test).

**Fig. S11** AR8 promotes Choy Sum growth in urban farm conditions. (a) Representative images of Choy Sum plants grown in coco peat with water (mock control) or AR8 conidia (10^6^ spores per coco peat) in outdoor field conditions. Choy Sum seeds placed on the surface of coco peat bungs (ensuring 1 seedling per bung per pot) were inoculated with water or with AR8 conidial suspension. (b) Shoot fresh weight at 28 dpi of Choy Sum seedlings inoculated with or without AR8. The boxes reveal the first quartile, median and third quartile; whereas the whiskers indicate the minimum and maximum values. Asterisks (***) represent significantly different means compared to the mock control at *P* < 0.001 (t-test).

## Supporting information

**Fig. S1** Fungal isolates from Arabidopsis rhizosphere modulate Choy Sum growth.

**Fig. S2** AR8 promotes plant growth and fertility under soil conditions.

**Fig. S3** AR8 promotes growth in Brassicaceae plants under soil conditions.

**Fig. S4** AR8 promotes growth in cereal crops under soil conditions.

**Fig. S5** AR8 solubilizes inorganic Pi sources in Pikovskaya broth.

**Fig. S6** Overview of metabolic changes in Choy Sum upon AR8 inoculation.

**Fig. S7** Repertoire of primary metabolites (sugars and amino acids) in Choy Sum upon AR8 inoculation.

**Fig. S8** Repertoire of phenylpropanoid (hydroxycinnamic acids) in Choy Sum upon AR8 inoculation.

**Fig. S9** Impact of hydroxycinnamic acids on the growth of Choy Sum.

**Fig. S10** *trans-*Cinnamic acid displays growth-promoting activity in Choy Sum plants in a dosage-dependent manner.

**Fig. S11** AR8 promotes Choy Sum growth in urban farm conditions.

**Methods S1** Fungal molecular identification and growth conditions.

**Methods S2** Pi solubilizing activity assay.

**Methods S3** Phenotypic characterization of the effect of hydroxycinnamic acids on Choy Sum.

**Methods S4** Metabolite extraction

**Methods S5** Detailed targeted analysis using GC/LC-MS technology for metabolites quantification.

**Methods S6** GC-MS data processing.

**Methods S7** Outdoor field trial or urban farm experiments.

**Dataset S1.** Normalized metabolites data from GC-MS and LC-MS analysis in shoot of Choy Sum upon water (mock control) and AR8 inoculation.

