## Supplemental Information for "Mycorrhizal symbiont provides growth benefits in host plants via phosphate and phenylpropanoid metabolism"

**BioRxiv Supporting Information file**

The following Supporting Information is available for this article:

*Supplementary Figures*

**Fig. S1** Fungal isolates from Arabidopsis rhizosphere modulate Choy Sum growth.

**Fig. S2** AR8 promotes plant growth and fertility under soil conditions.

**Fig. S3** AR8 promotes growth in Brassicaceae plants under soil conditions.

**Fig. S4** AR8 promotes growth in cereal crops under soil conditions.

**Fig. S5** AR8 solubilizes inorganic Pi sources in Pikovskaya broth.

**Fig. S6** Overview of metabolic changes in Choy Sum upon AR8 inoculation.

**Fig. S7** Repertoire of primary metabolites (sugars and amino acids) in Choy Sum upon AR8 inoculation.

**Fig. S8** Repertoire of phenylpropanoid (hydroxycinnamic acids) in Choy Sum upon AR8 inoculation.

**Fig. S9** Impact of hydroxycinnamic acids on the growth of Choy Sum.

**Fig. S10** *trans*-Cinnamic acid displays growth-promoting activity in Choy Sum plants in a dosage-dependent manner.

**Fig. S11** AR8 promotes Choy Sum growth in urban farm conditions.

*Supplementary Methods*

**Methods S1** Fungal molecular identification and growth conditions.

**Methods S2** Pi solubilizing activity assay.

**Methods S3** Phenotypic characterization of hydroxycinnamic acids on Choy Sum.

**Methods S4** Metabolite extraction

**Methods S5** Detailed targeted analysis using GC/LC-MS technology for metabolites quantification.

**Methods S6** GC-MS data processing.

**Methods S7** Field/Urban farm experiments.

*Supplementary Dataset*

**Dataset S1.** Normalized metabolomics data from GC-MS and LC-MS analysis in shoot of Choy Sum upon water (mock control) or AR8 conidial inoculation.

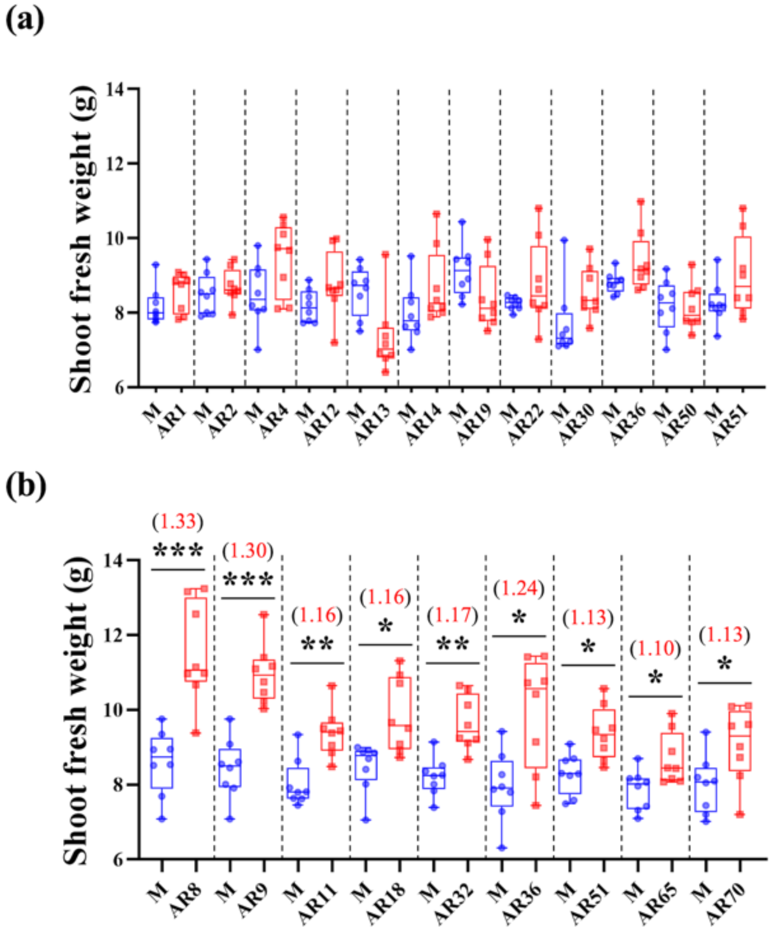


**Fig. S1** Fungal isolates from Arabidopsis rhizosphere modulate Choy Sum growth. (a-b) Shoot fresh weight of Choy Sum plants inoculated with non-PGPF isolates (AR1, AR2, AR4, AR12, AR13, AR14, AR19, AR22, AR30, AR36, AR50, and AR51) (a) or PGPF isolates (AR8, AR9, AR11, AR18, AR32, AR36, AR51, AR65, and AR70) (b) in soil conditions. Choy Sum seedlings (n=8 plants per experiment; three replicates of the experiment) inoculated with water (mock control) or conidia ($10^6$ spores in total) from the indicated fungal isolate in each instance, and shoot fresh weight measured at 21 days post-inoculation (dpi). M refers to mock control inoculated with water. Numbers in (b) represent the fold change in shoot fresh weight between fungal inoculation and mock control. The boxes reveal the first quartile, median and third quartile; the whiskers indicate the minimum and maximum values. Asterisks represent significantly different means compared to the mock control at *$P < 0.05$, **$P < 0.01$, ***$P < 0.001$ (Student's t-test).

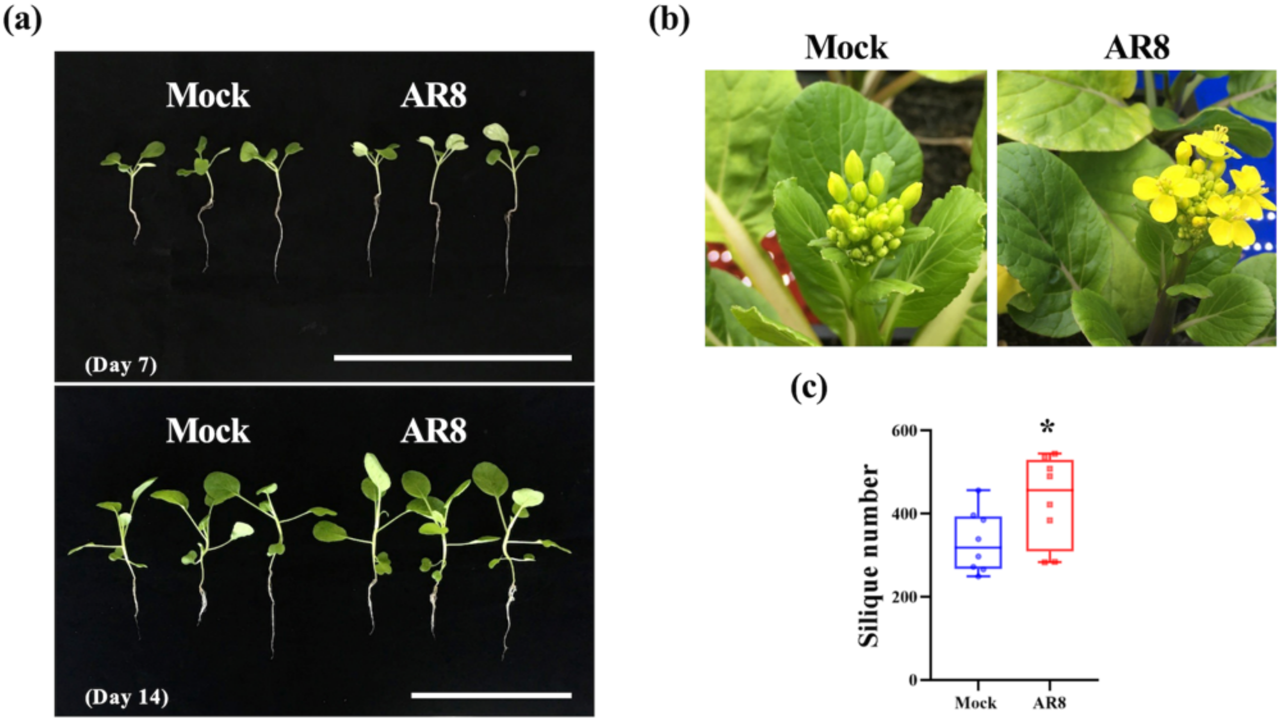


**Fig. S2** AR8 promotes plant growth and fertility under soil conditions. (a) Representative images of Choy Sum seedlings grown in soil with water (mock control) or AR8 conidia ($10^6$ spores) for 7 and 14 days. Scale bar, 10 cm. (b) Representative images of floral initiation time point in mock control or AR8-inoculated Choy Sum plants. Images shown are at 28 dpi. (c) Silique number produced by Choy Sum grown in soil measured at 49 dpi. Data presented (mean ± S.E) was derived from 3 independent replicates of the experiment. The boxes reveal the first quartile, median and third quartile; the whiskers indicate the minimum and maximum values. Asterisks (*) represent significantly different means compared to the mock control at $P < 0.05$ (t-test).

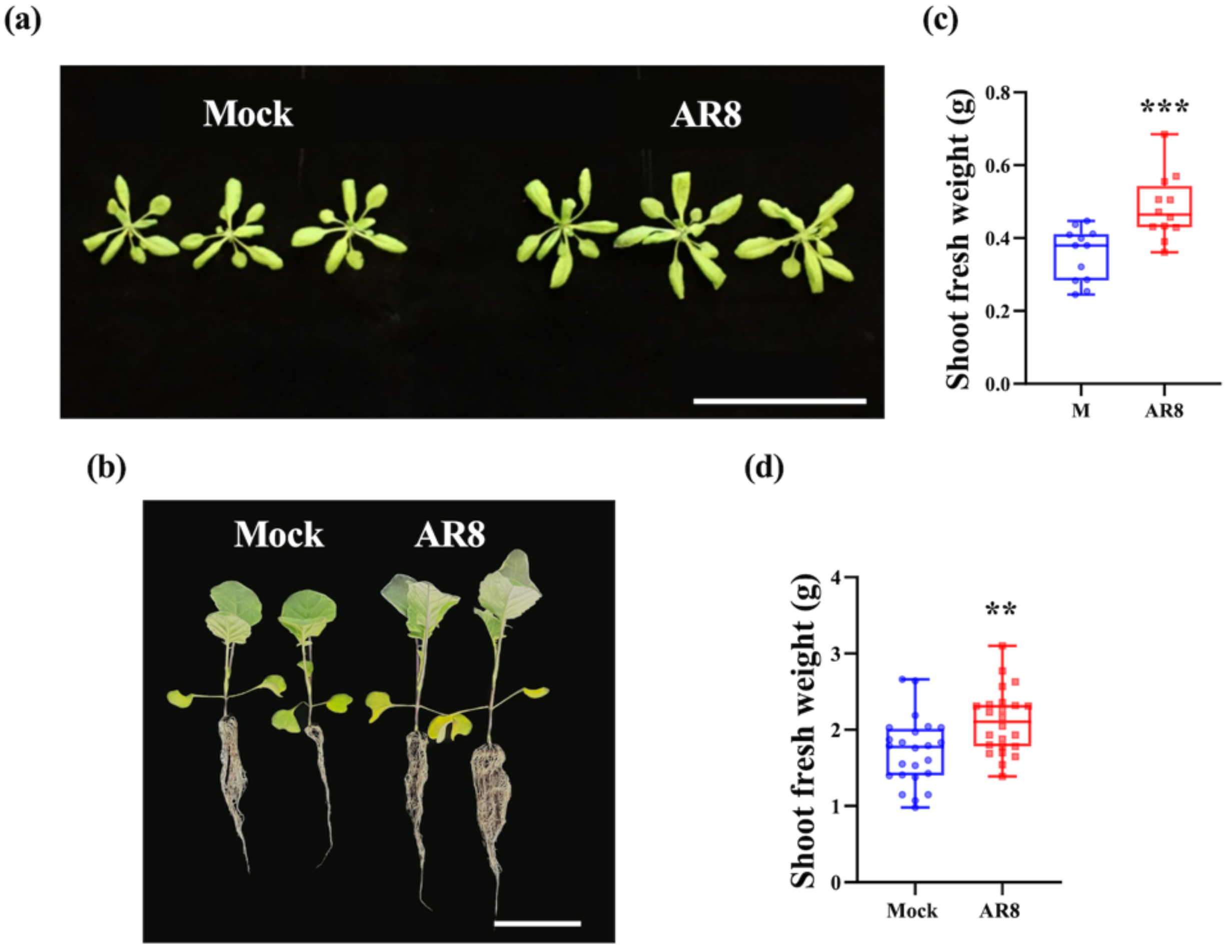


**Fig. S3** AR8 promotes growth in Brassicaceae plants under soil conditions. (a-b) Representative images of *Arabidopsis thaliana* Col-0 (a) and Kailan (b) grown in soil with water (mock control) or AR8 conidia ($10^6$ spores) for 21 days. Scale bars, 10 cm. (c-d) Shoot fresh weight of *Arabidopsis thaliana* Col-0 (c) and Kailan (d) grown in soil inoculated with water or AR8 conidia (n=12 and 24 plants per experiment, respectively; three replicates of the experiment). The boxes reveal the first quartile, median and third quartile; the whiskers indicate the minimum and maximum values. Asterisks represent significantly different means compared to the mock control at $^{**}P < 0.01$, $^{***}P < 0.001$ (t-test).

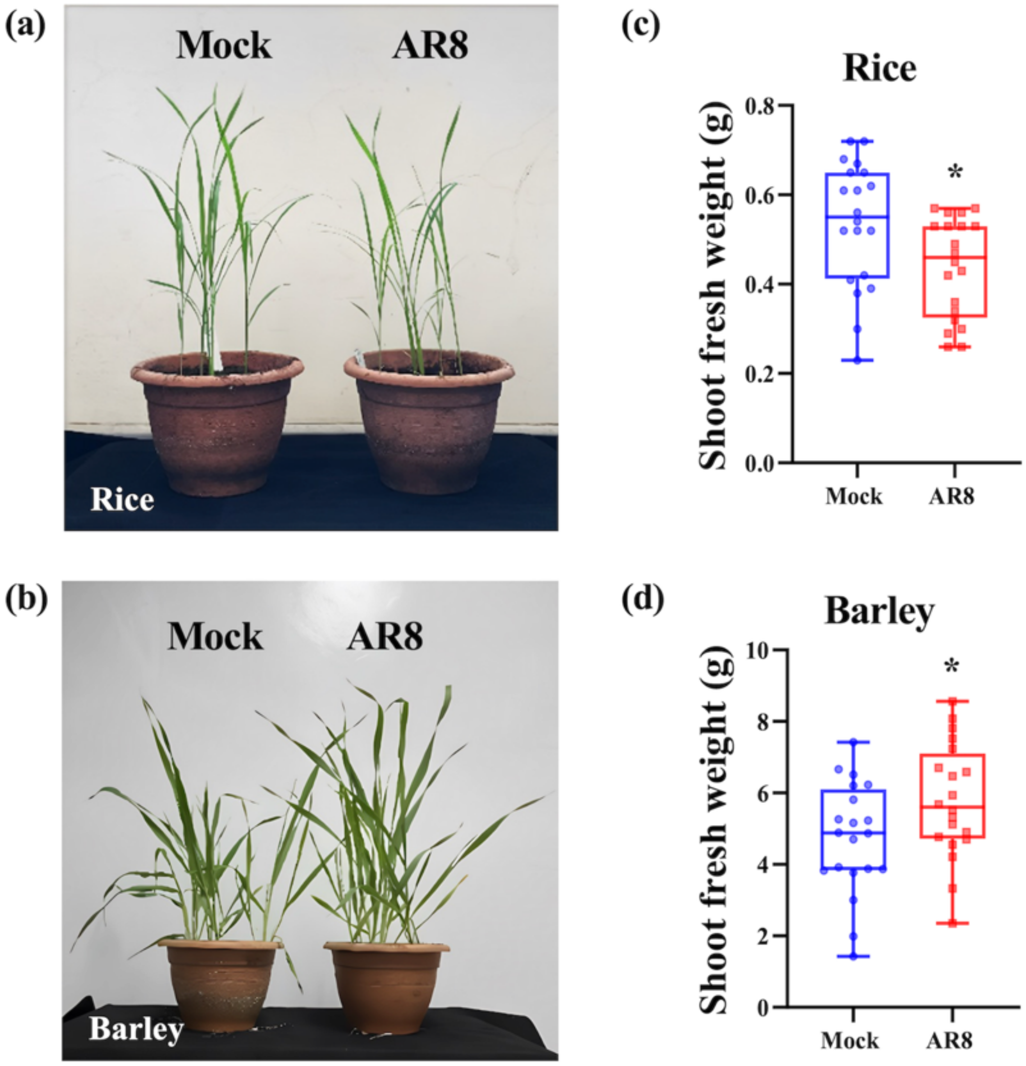


**Fig. S4** AR8 promotes growth in cereal crops under soil conditions. (a-b) Representative images of rice cultivar CO39 (a) and barley cultivar Express (b) grown in soil with water (mock control) or AR8 conidia ($10^6$ spores) for 21 days. (c-d) Shoot fresh weight of rice cultivar CO39 (c) and barley cultivar Express (d) grown in soil inoculated with water or AR8 conidia (n=20 plants per experiment, respectively; three replicates of the experiment). The boxes reveal the first quartile, median and third quartile; the whiskers indicate the minimum and maximum values. Asterisks represent significantly different means compared to the mock control at $*P < 0.05$ (t-test).

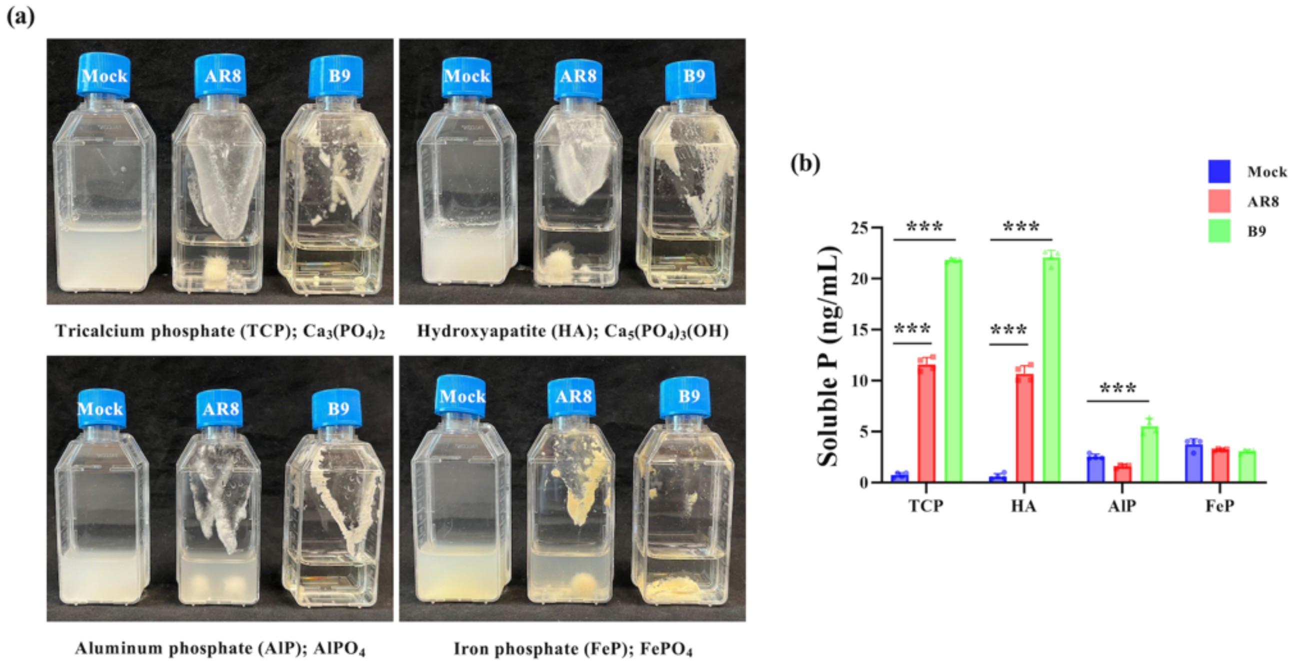


**Fig. S5** AR8 solubilizes inorganic Pi sources in Pikovskaya broth. (a) Pi solubilizing capability of AR8 and B9 in different inorganic Pi sources (tricalcium phosphate, hydroxyapatite, aluminum phosphate, and iron phosphate) after 7 days. (b) Soluble Pi concentration produced by AR8 and B9 in in Pikovskaya broth with different inorganic Pi sources. Mycelial plugs (AR8 and B9) were inoculated to Pikovskaya broth with different inorganic Pi sources for 7 days and the soluble Pi concentration was measured using phosphomolybdenum spectrophotometry. Data presented (mean ± S.E) was derived from 3 independent replicates of the experiment. Asterisks represent significant differences compared to the mock control at ***$P < 0.001$ (t-test).

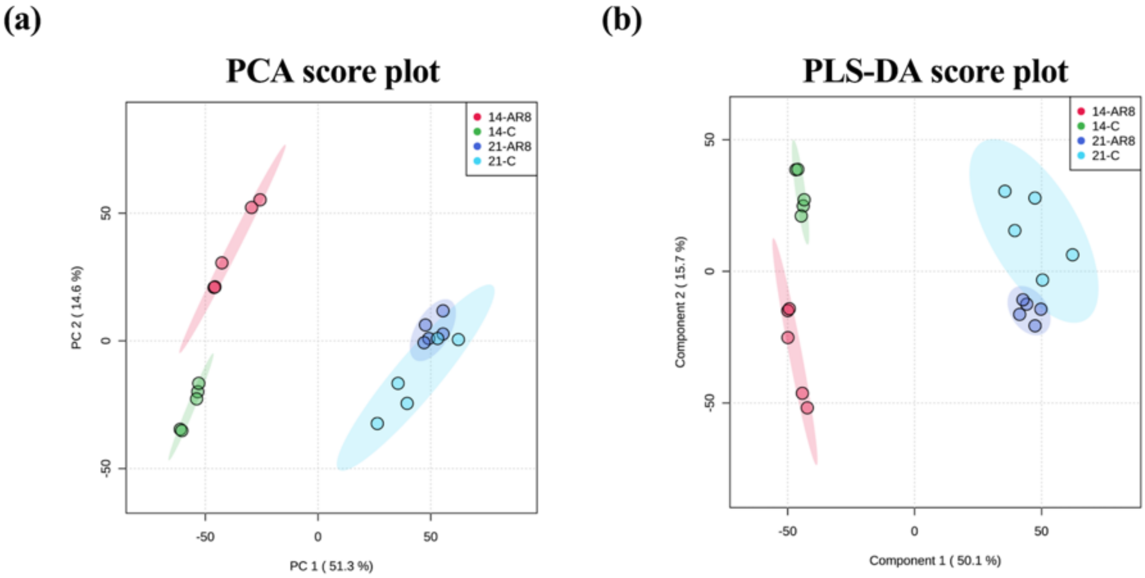


**Fig. S6** Overview of metabolic changes in Choy Sum upon AR8 inoculation. (a-b) The PCA (a) and PLS-DA (b) score plots for shoots metabolome of Choy Sum plants inoculated with water or AR8 conidia ($10^6$ spores) at 14 and 21 dpi as captured by GC-EI/TOF-MS. The colored ellipses represent 95% confidence regions for each group.

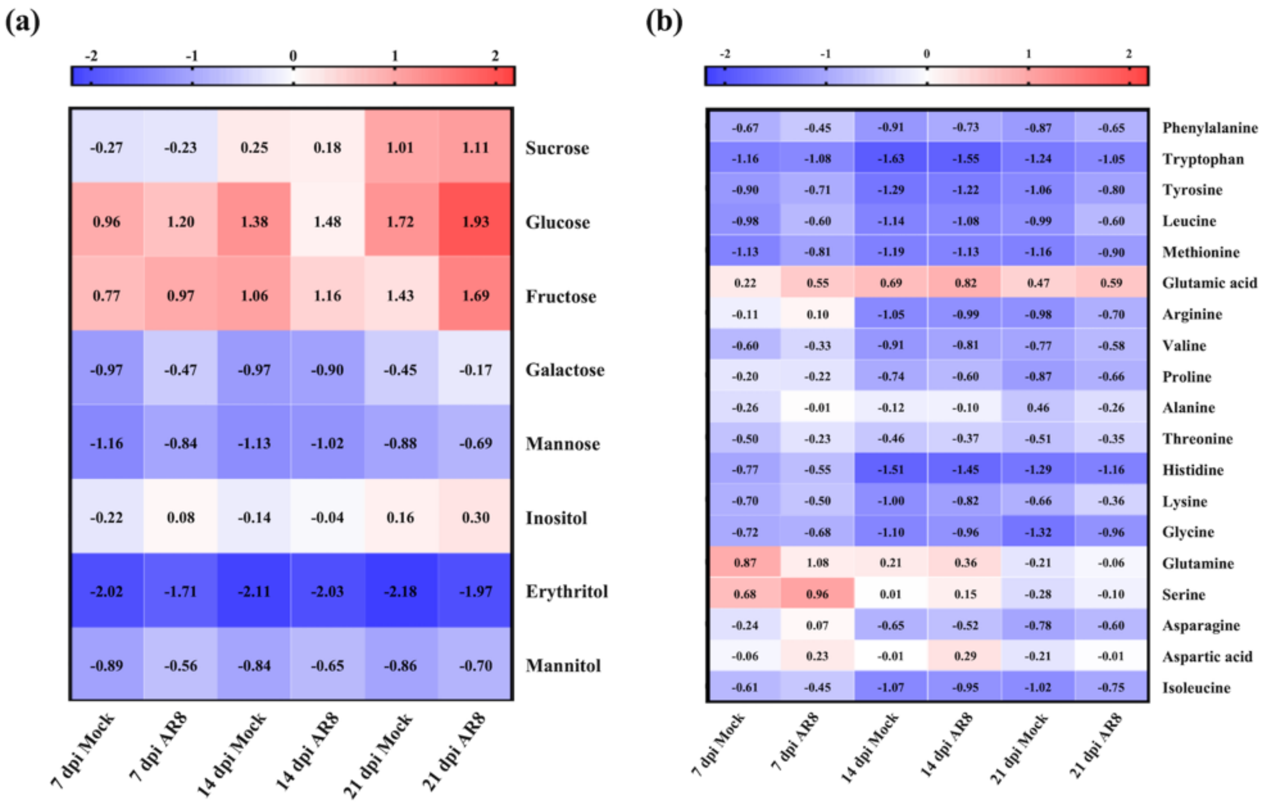


**Fig. S7** Repertoire of primary metabolites (sugars and amino acids) accumulating in Choy Sum during beneficial association with AR8. (a-b) Heatmap showing quantitative and qualitative changes in concentration of sugars (a) and amino acids (b) in Choy Sum shoots at three growth stages (microgreen, seedling, and adult) upon water or AR8 conidial inoculation were performed by targeted analysis. The heatmap was generated through log transformation and colored by concentration (μg/g), e.g., 10 becomes 1, highlighted in blue (low) or red (high) via GraphPad Prism 8, respectively. Numbers represent the average concentration of corresponding metabolites derived from five biological replicates.

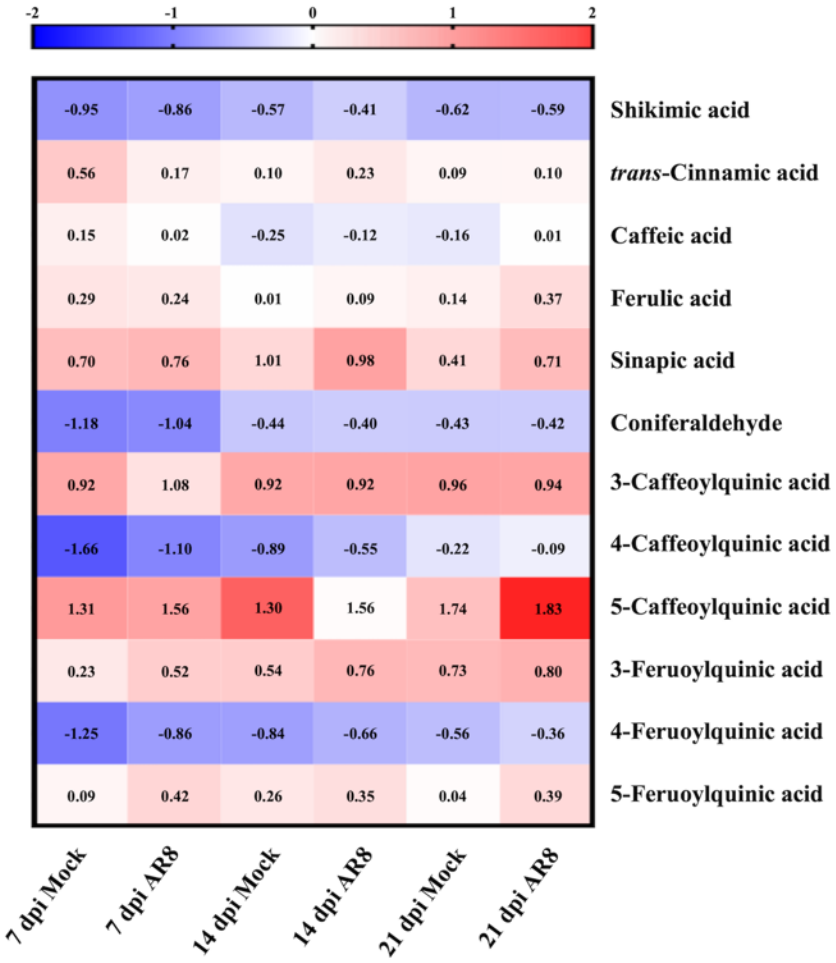


**Fig. S8** Repertoire of phenylpropanoids (hydroxycinnamic acids) accumulating in Choy Sum during beneficial association with AR8. Heatmap showing quantitative and qualitative changes in abundance of hydroxycinnamic acids and their derivatives in Choy Sum shoots at three growth stages (microgreen, seedling, and adult) upon water or AR8 inoculation were performed by targeted analysis. The heatmap was generated through log transformation and colored by concentration (μg/g), e.g., 10 becomes 1, highlighted in blue (low) or red (high) via GraphPad Prism 8, respectively. Numbers represent the average concentration of corresponding metabolites abundance of five biological replicates.

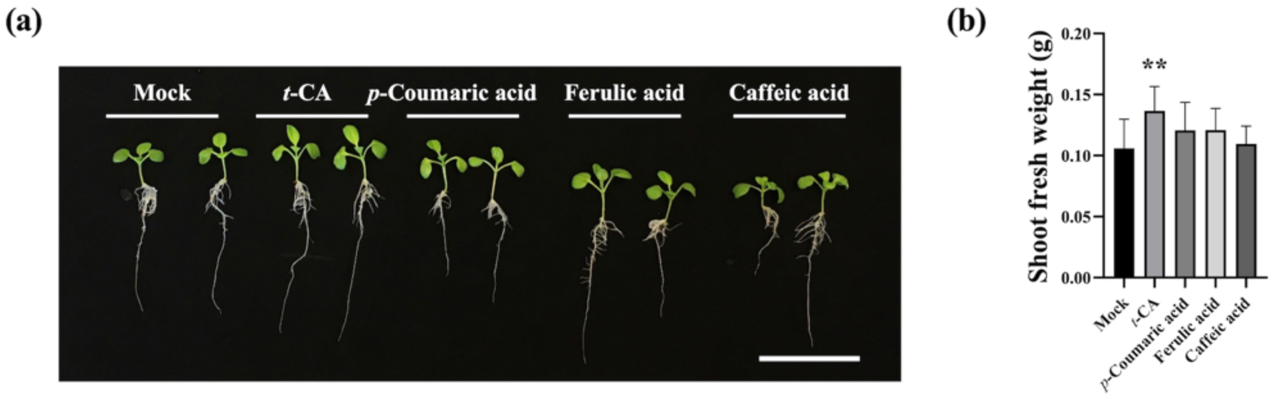


**Fig. S9** Impact of specific hydroxycinnamic acids on the growth of Choy Sum. (a) Representative images of Choy Sum seedlings grown in MS medium containing *t*-CA (2.5 μM), *p*-coumaric acid (60 μM), ferulic acid (20 μM), and caffeic acid (20 μM) for 7 days. Scale bar, 5 cm. (b) Shoot fresh weight of Choy Sum grown on control and hydroxycinnamic acids-supplemented (*t*-CA, *p*-coumaric acid, ferulic acid, and caffeic acid) medium for 7 days (n=10 plants per experiment). Data presented (mean ± S.E) were derived from 3 independent replicates of the experiment. Asterisks represent significant differences compared to the mock control at $^{**}P < 0.01$ (t-test).

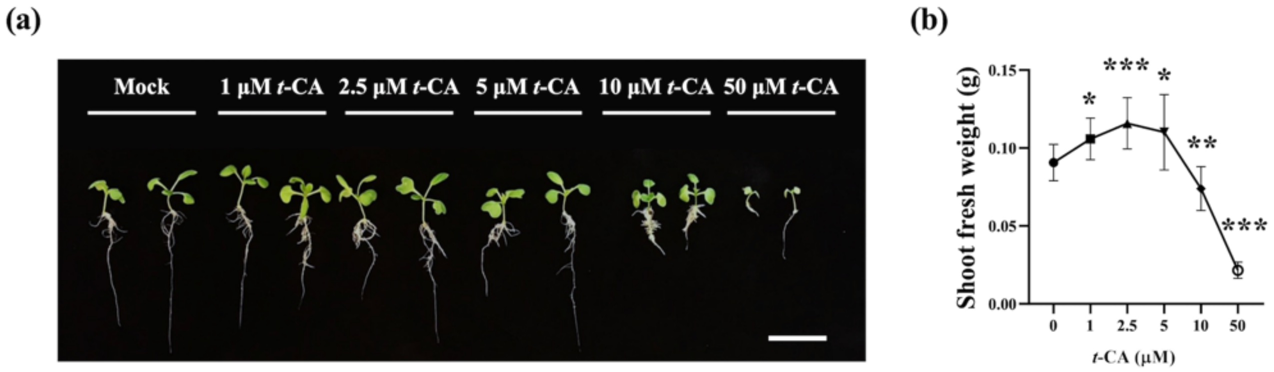


**Fig. S10** *trans*-Cinnamic acid displays growth-promoting activity in Choy Sum plants in a dosage-dependent manner. (a) Representative images of Choy Sum seedlings grown in MS medium containing the denoted amounts of *t*-CA (0, 1, 2.5, 5. 10, and 50 μM) for 7 days. Scale bar, 5 cm. (b) Shoot fresh weight of Choy Sum grown on uninoculated control or *t*-CA-supplemented medium for 7 days (n=10 plants per experiment). Data presented (mean ± S.E) were derived from 3 independent replicates of the experiment. Asterisks represent significantly different means compared to the mock control at *$P < 0.05$, **$P < 0.01$, ***$P < 0.001$ (t-test).

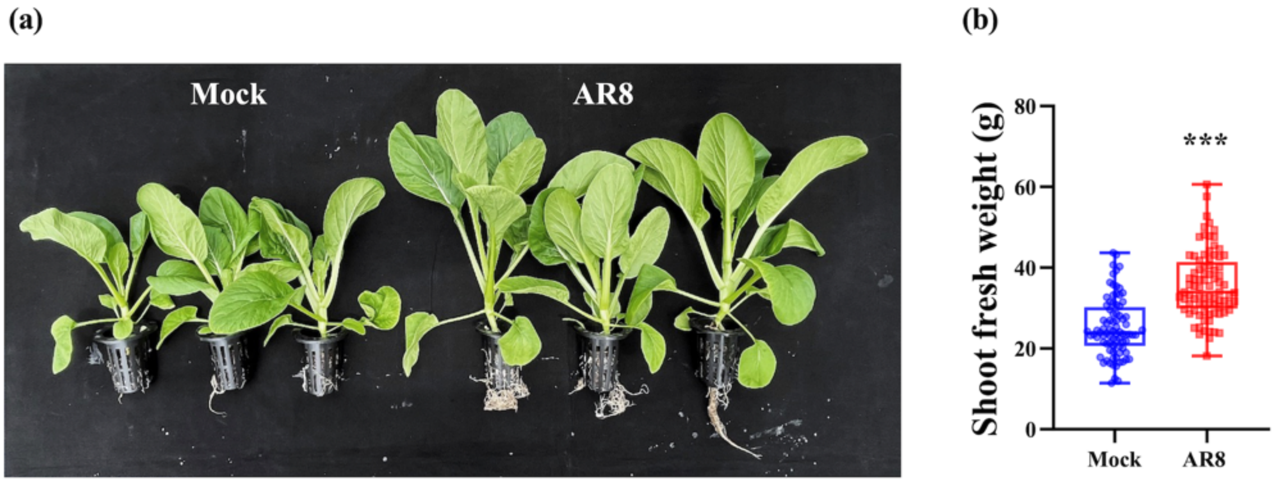


**Fig. S11** AR8 promotes Choy Sum growth in urban farm conditions. (a) Representative images of Choy Sum plants grown in coco peat with water (mock control) or AR8 conidia ($10^6$ spores per coco peat) in outdoor field conditions. Choy Sum seeds placed on the surface of coco peat bungs (ensuring 1 seedling per bung per pot) were inoculated with water or with AR8 conidial suspension. (b) Shoot fresh weight at 28 dpi of Choy Sum seedlings inoculated with or without AR8. The boxes reveal the first quartile, median and third quartile; whereas the whiskers indicate the minimum and maximum values. Asterisks (***) represent significantly different means compared to the mock control at $P < 0.001$ (t-test).

### ***Supplementary Methods***

**Methods S1** Fungal molecular identification and growth conditions

Healthy *A. thaliana* Col-0 roots were collected and fungal microbiota extracted as described previously (Gu et al., 2023). Briefly, root tissue samples were ground in phosphate buffered saline and the homogenate was diluted serially and plated on Prune juice agar (PA) medium containing antibiotics and incubated at 28°C for 2-3 days. The individual fungal colonies were selected and cultured on PA medium with 50 mm filter-paper discs twice for purification, and storage. To identify the fungal isolates, genomic DNA from mycelia grown in Complete (CM) medium was extracted using the MasterPure$^{TM}$ Yeast DNA purification kit (Lucigen Corporation, Middleton, Wisconsin, USA) and 10 ng of fungal DNA template used for PCR amplification with specific sets of primers that target the internal transcribed spacer (ITS) region (White et al., 1990). PCR products were sequenced and DNA barcoding performed using the NCBI BLAST database (https://blast.ncbi.nlm.nih.gov/Blast.cgi), assigning the genus and/or species identity of our fungal isolates by utilizing the highest identity and/or similarity scores.

**Methods S2** Pi solubilizing activity assay

Mycelial plugs (5 mm) with or without AR8 were inoculated to the modified Pikovskaya broth containing insoluble inorganic Pi (tricalcium phosphate, iron phosphate, aluminum phosphate, and hydroxyapatite) (Nautiyal, 1999). *Penicillium citrinum* isolate B9, which has been characterized for Pi solubilization activity was used as a positive control for comparison (Gu et al., 2023). Samples were incubated at 28°C, 180 rpm on an orbital shaker for 7 days. The cell-free culture medium was collected by filtering through 4 layers of Miracloth and centrifuged at 4000 rpm for 5 min. The soluble phosphorus content was determined using an Ultraspec 8000 spectrophotometer (Scimed Asia) at 430 nm and the amount of soluble phosphate produced was calculated using a standard curve of $KH_2PO_4$ prepared in Pikovskaya broth.

**Methods S3** Phenotypic characterization of the effect of hydroxycinnamic acids on Choy Sum

To examine the effect of phenylpropanoids on plant growth, 4-day-old Choy Sum seedlings were transferred to the sucrose-free MS-agar medium supplemented with corresponding metabolites in PhytaTray II (Sigma-Aldrich) for 7 days. The methodology and concentrations of phenylpropanoids used in the study were according to a previous publication (Kurepa et al., 2018). *t*-CA was further used in the dose-dependent treatment and co-culture assays with AR8 on Choy Sum seedlings. AR8 mycelial plugs (5 mm each) were inoculated in the PhytaTray with or without t-CA for 7 days and 4-day-old Choy Sum seedlings were then transferred to the PhytaTray in the presence of t-CA, AR8, or t-CA with AR8, and cultivated for 7 days under either incandescent tube light or LED light conditions. Shoot fresh weight was measured to study the complementary association between AR8 and *t*-CA.

**Methods S4** Metabolite extraction

Metabolite analysis was carried out using previously established protocols (Zou et al., 2021). Briefly, freeze-dried Choy Sum powder (50 mg) was extracted thrice by 1 ml of precooled 80% methanol (v/v) under sonication in an ice bath for 15 min and then centrifuged for 10 min at 14,000 rpm at 5°C. For amino acids, plant extracts were further vacuum-dried and reconstituted in 3 ml of 70% acetonitrile (v/v) for analysis. For derivatization, plant extracts were vacuumed-dried and derivatized with methoxyamine (5 mg/ml in pyridine) for 2 hours at 60 °C. The derivatives were subsequently trimethylsilylated with 70 µl of N-methyl-N-(trimethylsilyl) trifluoroacetamide for 1 hour at 37°C.

**Methods S5** Detailed targeted analysis using GC/LC-MS technology for metabolites quantification

#### Quantification of sugar content in Choy Sum

The quantitative sugar analysis was performed by gas chromatography coupled to a 7000B Triple Quadrupole mass detector (Agilent Technologies, Palo Alto, CA) equipped with a chemical ionization source onto A J&W HP-5MS-23 column (60 m length, 0.25 mm inner diameter, 0.25 µm film thickness, Agilent). Helium was used as carrier gas and the flow rate was 2 mL/min. Injection volume was 1 µL in split-less mode. The GC oven temperature was held at 120°C for 1 minute, then increased to 200°C at 25°C/min, then increased to 217°C at 1.5°C/min, and eventually increased to 300°C at 25°C/min and held for 5 minutes. Mass spectra were acquired in positive multiple reaction monitoring (MRM) mode with following parameters: methane as reactant gas, and gas flow was set at 20%, ion source temperature was at 250°C.

**Quantification of amino acid content in Choy Sum**

Amino acid analysis was performed in an Agilent 1200 HPLC system (Agilent, Germany) coupled to a 6410 Triple Quadrupole (Agilent, USA) mass spectrometer equipped with an electrospray ionization source. The separation of compounds was achieved on an Acquity UPLC BEH amide column (100 mm length, 2.1mm inner diameter, 1.7 µm particle size, 100 Å, Waters). The autosampler and column were maintained at 4 °C and 35 °C, respectively. For analysis, plant samples were injected with 5 µL. Separation was performed at a constant flow rate of 0.5 mL/min, in a gradient of solvent A (30% acetonitrile containing 10 mM ammonium formate with 0.1% formic acid) and B (95% acetonitrile containing 10 mM ammonium formate with 0.1% formic acid). The gradient program was: The gradient program was: 100% solvent B (0-1 minute); 100-92% solvent B (1-2 minute); 92-85% solvent B (2-10 minute); 85-60% solvent B (10-12 minute); 60-40% solvent B (12-14 minute); 40-15% solvent B (14-15 minute); 15% solvent B (15-19 minute); re-equilibration from 15-100% solvent B (19-20 minute); and reconditioned with 100 % solvent B (20-35 minute). Electrospray ionization source was performed in negative ion mode with the following source parameters: drying gas ($N_2$) temperature of 350°C with a flow of 12 L/min, nebulizer gas pressure of 30 psi, sheath gas temperature of 400 °C with a flow of 11 L/min and capillary voltage of 3500 V. Mass spectra were acquired in the MRM mode.

**Quantification of phenylpropanoid content in Choy Sum**

Phenylpropanoid analysis was performed in an Agilent 1290 Infinity II coupled to an Agilent 6495 Series Triple Quadrupole LCMS (Agilent, USA) with a Jet Stream electrospray ionization source ion source. The separation of compounds was achieved on a Kinetex C18 column (100 mm length, 2.1mm inner diameter, 1.7 µm particle size, 100 Å, Phenomenex). The autosampler and column were maintained at 4°C and 35°C, respectively. For analysis, plant samples were injected with 5 µL. Separation was performed at a constant flow rate of 0.3 mL/min, in a gradient of solvent A (water acidified with 0.1% formic acid) and B (acetonitrile acidified with 0.1% formic acid). The gradient program was 2-20% solvent B (0-5 minute), 20-30% solvent B (5-12 minute); 30%-100% solvent B (12-12.1 minute); 100% solvent B (12.1-15 minute); re-equilibration from 100-2% solvent B (15-15.1 minute); and reconditioned with 2 % solvent B (15.1-19 minute). Electrospray ionization source was performed in negative ion mode with the following source parameters: drying gas (N2) temperature of 280°C with a flow of 12 l/min, nebulizer gas pressure of 40 psi, sheath gas temperature of 350°C with a flow of 12 l/min and capillary voltage of 3000 V. Mass spectra were acquired in the MRM mode.

**Methods S6** GC-MS data processing

The raw data acquired from GC-MS were first transformed to mzData by the MassHunter Qualitative Analysis software (Agilent). The results were then exported to MZmine software for peak alignment. All aligned features were normalized to the abundance of internal standard (FMOC-glycine). Subsequently, the normalized features were screened using the criteria of relative standard deviation (RSD) and detection frequency (DF) for the same feature in the same group. Notably, the features higher than 30% RSDs or lower than 80% DFs were excluded from the following statistical analysis. The features with variable importance in projection (VIP) scores above 1 in the OPLS-DA model and p-values lower than 0.05 in the Kruskal-Wallis test were considered to be statistically significant and selected to identify their corresponding metabolites.

The metabolites in GC-MS analysis were identified on the basis of mass spectra and accurate mass of fragments by searching the National Institute of Standards and Technology (NIST) 11 library. Retention time index was applied to confirm the identification of metabolites in GC-MS analysis. All mass features were normalized to the sample fresh weight and internal standard. For quantification, all features were evaluated for the best specific and quantitative representation of observed targets.

**Methods S7** Outdoor field trial or urban farm experiments

The field trials were conducted at a local outdoor urban farm in Singapore. To quantify the AR8 plant growth-promoting effect under natural outdoor conditions, fungal inoculation was performed directly on Choy Sum seeds at the time of sowing. About 1-2 (unsterilized) seeds were sown on the surface of coco peat (114 coco peat bungs per group in total for each trial, with 4 trials or repeats in total) and inoculated with AR8 conidial suspension ($1x10^6$ spores). Distilled water was used as a mock control with the same inoculation method. The irrigation system was kept on for 24 hours to keep water flowing through the base of the coco peat bungs placed in the perforated plastic pots. Seed germination was checked at 7 dpi. Additional seedlings (per pot; if any) were removed to retain and maintain only one properly germinated seed/seedling in each coco peat to prevent the shade avoidance syndrome. Shoot fresh weight of Choy Sum plants was measured at 28 dpi.
